# mm2-ivh: simple and precise overlap detection in alpha satellite HORs with interval hashing

**DOI:** 10.1101/2025.07.11.664273

**Authors:** Hajime Suzuki, Masahiro Sugawa, Yoshitaka Sakamoto, Yuichi Shiraishi

## Abstract

**Summary:** We propose a new algorithm, “interval hashing,” which distinguishes identical k-mers arising from different repeat sequences, particularly in complex repeat arrays such as alpha satellite HORs. We implement this algorithm as a fork of minimap2, named mm2-ivh. In local assembly of alpha satellite HORs, mm2-ivh accurately reconstructs more haplotypes than assemblers using standard minimizers.

**Availability:** mm2-ivh is available under the MIT license at https://github.com/ocxtal/mm2-ivh and runs on common Unix-compatible systems.

## Introduction

Resolving repeat regions in genome assembly has been a persistent challenge. Recent advances in long-read sequencing technologies, including PacBio’s HiFi sequencing (Wenger *et al*. 2019) and Oxford Nanopore Technologies’ (ONT) ultra-long (UL) read protocol (Jain *et al*. 2018), have significantly improved the ability to resolve complex repeat regions. HiFi reads, which are approximately 10–20 kilobase pairs (kbp) long and have over 99.8% accuracy, enable the precise detection of minor variations between repeat elements (Dvorkina, Bzikadze and Pevzner 2020). UL reads, which exceed 100 kbp in length, enable the resolution of long and complex repeat element arrangements that were previously intractable (Miga *et al*. 2020; Logsdon *et al*. 2021; Nurk *et al*. 2022). Assemblers such as Hifiasm (Cheng *et al*. 2021) and Verkko (Rautiainen *et al*. 2023) use HiFi reads to build accurate assembly graphs and UL reads to untangle these graphs into linear contigs, enabling the construction of nearly complete telomere-to-telomere sequences for a considerable number of chromosomes (Yang *et al*. 2023; Logsdon *et al*. 2024).

Nevertheless, accurately assembling long and highly homologous repeat regions remains challenging. For instance, the centromeric alpha satellite consists of 171 bp monomers that repeat over several megabases in a higher-order repeat (HOR) structure (Wu and Manuelidis 1980; Willard and Waye 1987; Thakur, Packiaraj and Henikoff 2021; Logsdon and Eichler 2022). Several methods have been developed to assemble and validate alpha satellite HORs using “unique k-mers,” observed only at specific locations within HOR regions, enabling the accurate reconstruction of HOR structures across several chromosomes (Bzikadze and Pevzner 2020; Miga *et al*. 2020; Logsdon *et al*. 2021; Bzikadze, Mikheenko and Pevzner 2022; Dishuck *et al*. 2023; Chakravarty, Logsdon and Lonardi 2025). In long-read mapping within HOR regions and comparisons between HOR regions, methods that adaptively adjust the length of seeds based on uniqueness have also proven effective (Jain *et al*. 2022; Bzikadze and Pevzner 2023; Zhang *et al*. 2025). However, a broadly applicable method for overlap detection in diverse alpha satellite HORs has yet to be developed. Even state-of-the-art assemblers often produce misassemblies or inconsistencies among tools in alpha satellite HORs (Logsdon *et al*. 2025).

To address the challenges of assembling alpha satellite HORs, we propose a simple yet effective algorithm to improve seeding in periodic repeats. This algorithm, interval hashing, encodes the distances between occurrences of identical k-mers or minimizers (Roberts *et al*. 2004) into hash values, enabling the distinction of identical k-mers based on their surrounding repeat contexts. We implemented the algorithm on top of minimap2 (Li 2018) and applied it in combination with miniasm (Li 2016) for the local assembly of alpha satellite HORs using high-quality human ONT UL reads. Minimap2 with interval hashing successfully reconstructed the full-length active HOR regions in 27 out of 46 haplotypes without macroscopic misassemblies, outperforming the original minimap2 and Hifiasm in ONT-only mode. The implemented tool, mm2-ivh, is freely available under the MIT license at https://github.com/ocxtal/mm2-ivh.

### Algorithm

The difficulty of assembling alpha satellite HORs arises from the low specificity of seeds, which is caused by the presence of identical k-mers in multiple monomers. To improve seed specificity and make the overlap detection easier in alpha satellite HORs, we leverage an intrinsic property of the regions. In alpha satellite HORs, certain k-mers appear only in specific monomers and generate characteristic, sometimes non-equidistant patterns of k-mer occurrence intervals. The presence of truncated or rare HOR units, or the insertion of unrelated sequences such as transposable elements, also leads to distinct occurrence interval patterns compared to regions without such events. Our algorithm, interval hashing, encodes the k-mer occurrence intervals into hash values, serving as signatures to distinguish identical k-mers by their surrounding contexts (Fig. 1(a)).

**Figure 1.**
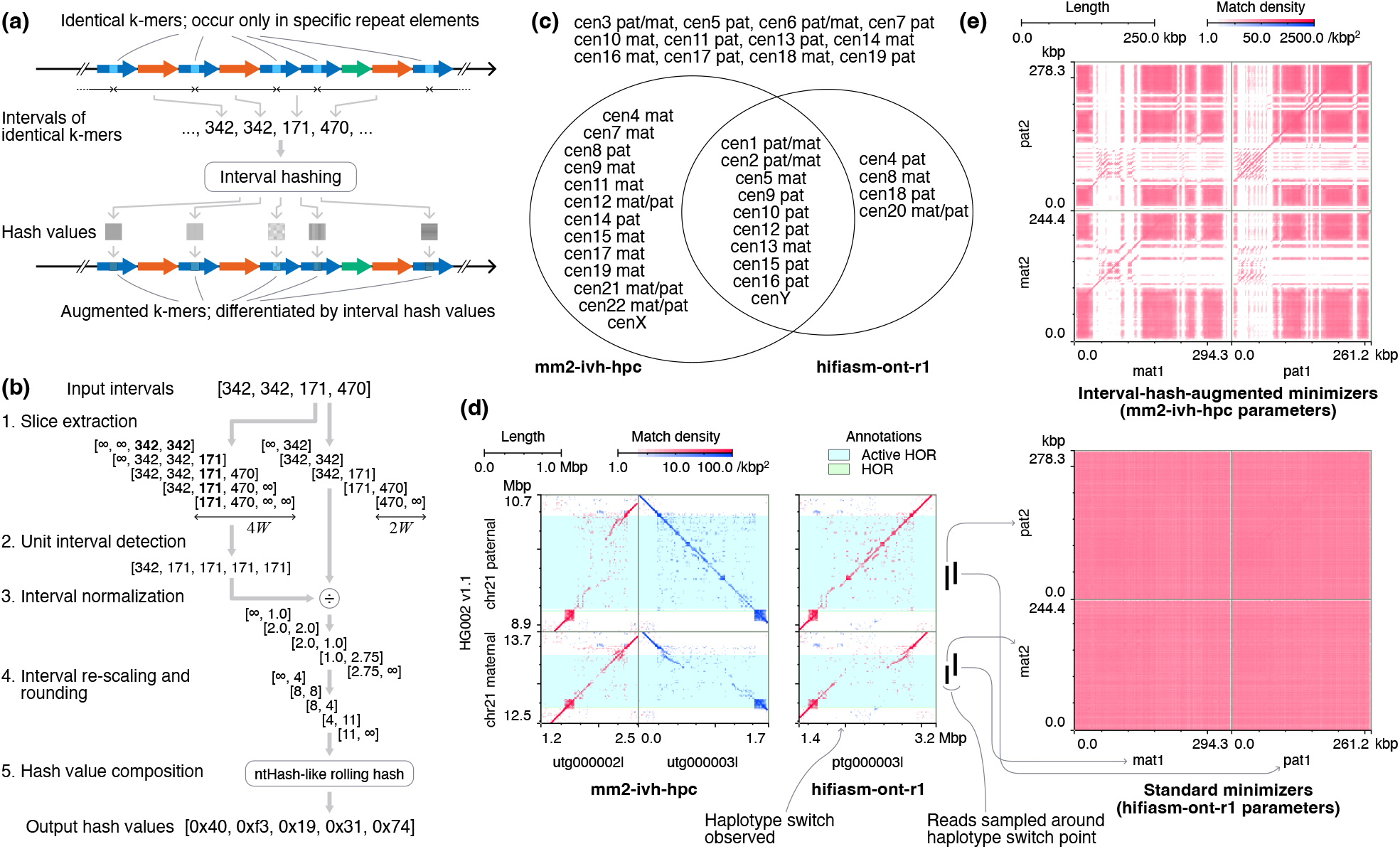
**(a)** Concept of interval hashing. **(b)** Flow of the five steps of the core of our algorithm (depicts *N* = 4 and *W* = 1). **(c)** Haplotypes in which the full length of active HOR regions is reconstructed into a single unitig. **(d)** Dotplots of the assembled unitigs of alpha satellite HORs of chromosome 21 against the corresponding regions of HG002 v1.1. Only minimizers with (*k, w*) = (19, 15) occurring less than or equal to 10 times among all alpha satellite HOR regions in HG002 v1.1 are used. Red and blue dots are forward and reverse-complement matches, respectively. **(e)** Dotplots among four reads across the haplotype switch point in hifiasm-ont-r1 assembly. Reads mat1/2 and pat1/2 were primarily mapped across the corresponding position on the HG002 v1.1 chromosome 21 maternal and paternal, respectively. The colors are the same as in the panel (d).

The algorithm takes a vector of *N* (*N* ≥ 0) intervals, derived from the *N* + 1 occurrences of identical k-mers, and produces a vector of *N* + 1 hash values corresponding to each occurrence. The core algorithm is defined by a wing length parameter *W* that indicates the number of k-mer occurrences to consider in each direction and consists of five steps (Fig. 1(b)). In the first step, two vectors of *N* + 1 slices are extracted from the input intervals, with slice lengths being 2*W* and 4*W*. The central 2*W* span of *i*-th 4*W*-long slice overlaps with *i*-th 2*W*-long slice. The out-of-bound positions are assigned interval values of infinity. Next, the “unit interval,” representing the length of a canonical repeat element, is calculated for each 4*W*-long slice by taking the minimum interval value. These unit intervals are then used to normalize the interval values in the corresponding 2*W*-long slices. The normalized intervals are rounded to the nearest integer after being multiplied by a resolution constant (fixed at 4.0 in our implementation) to ensure sufficient resolution for truncated repeat elements. Then, the integer intervals in each slice are individually mapped to partial hash values and then combined into a final hash value using a rolling hashing algorithm similar to ntHash (Mohamadi *et al*. 2016). Infinite interval values are mapped to a special partial hash value of zero to indicate the absence of interval information, ensuring that the hash value computed for an empty input interval vector derived from singleton k-mers is zero.

In practical implementations, it is necessary to match k-mers that are in a reverse-complement relationship. Tools such as minimap2 canonicalize k-mers by selecting the numerically smaller k-mer between the two strands. Interval hashing achieves consistent canonicalization by reversing the slices when the corresponding central k-mer is computed from the reverse-complement strand. Additionally, to prevent encoding intervals that span distant repeat contexts, it is practical to split the interval vector at points where the intervals exceed a certain threshold before applying interval hashing.

### Implementation

We implemented the interval hashing algorithm in a fork of minimap2 2.28, which applies interval hashing to all minimizers. Further implementation details are provided in Supplementary Section S1. The executable is named mm2-ivh and is compatible with minimap2 in terms of options and input/output formats. Based on preliminary experiments described in Supplementary Section S3, we added a preset option, `-x ivh-ava-ont-ul`, tuned for the latest UL reads with a mean quality value of 20 or higher. The preset uses interval-hash-augmented minimizers with parameters (*k, w, W*) = (19, 15, 3) and sets the maximum minimizer separation for splitting interval vectors to 20 kbp.

mm2-ivh is available at https://github.com/ocxtal/mm2-ivh under the MIT license, following the original minimap2. The program is implemented solely in C, requiring only a C compiler and make to build. It supports x86_64 and arm64 systems running Linux or macOS. The version used in our experiments is also available as an archive at 10.5281/zenodo.17143522.

## Results

We evaluated mm2-ivh for the local assembly of human alpha satellite HOR regions using the HG002 high-accuracy UL data published by ONT (Supplementary Section S4.1). After quality control, we obtained 89.0 Gbp of reads with a mean length of 116.9 kbp and a mean quality value of 24.64. The reads were then aligned to CHM13 v2.0 (Nurk *et al*. 2022), which consists of the autosomes and chromosome X of CHM13 and chromosome Y derived from HG002, using minimap2 with the `-x map-ont`preset. Reads mapped to HOR regions with 500 kbp margins were extracted for each chromosome (Supplementary Table S3). The extracted reads were assembled using the following three assemblers (Supplementary Section S4.2). The first assembler computes overlaps using mm2-ivh and outputs a pairwise alignment format (PAF) file, which is then filtered to remove overlaps with low match rates and supplied to miniasm to generate assembly graphs (denoted as mm2-ivh-hpc). The parameters for mm2-ivh and miniasm were set as `-x ivh-ava-ont-ul -H`(additionally enabling homopolymer compression) and `-h5000 -c2`(setting the maximum overhang length to 5 kbp and minimum coverage to 2), respectively. The filtering step allows only overlaps with more than 20% match columns relative to the alignment length. The second assembler, based on mm2-ivh-hpc, disables interval hashing and instead increases the minimizer length to *k* = 133 (Supplementary Section S2 for details on supporting long minimizers; denoted as mm2-k133-hpc). The third assembler is Hifiasm 0.25.0, run with `--ont -r1`options (one round of error correction; denoted as hifiasm-ont-r1). The `-r1`setting was chosen because it yielded the best results in our tests.

mm2-ivh-hpc reconstructed unitigs that fully covered the active HOR regions of the HG002 v1.1 reference assembly (Rautiainen *et al*. 2023) in 27 out of 46 haplotypes, while hifiasm-ont-r1 achieved this in 16 haplotypes (Supplementary Section S4.3, Supplementary Table S8). mm2-k133-hpc reconstructed the full length of active HOR regions only in chromosome Y. The number of active HOR regions that could be reconstructed into a single unitig by both mm2-ivh-hpc and hifiasm-ont-r1 was 11, while 16 were reconstructed only by mm2-ivh-hpc, 5 only by hifiasm-ont-r1, and 14 were not reconstructed by either (Fig. 1(c)). The effectiveness of interval hashing is clearly demonstrated in some HOR arrays. For instance, for the active HOR region in chromosome 21, mm2-ivh-hpc was able to assemble the full length for both haplotypes without haplotype mixing, whereas hifiasm-ont-r1 introduced a haplotype switch in the middle (Fig. 1(d), Supplementary Section S4.4). The dotplot of four reads mapped across the haplotype switch position (Supplementary Table S9) showed that interval hashing clearly distinguished haplotypes, while standard minimizers obscured them with dense matches (Fig. 1(e), Supplementary Fig. S5).

Execution time and memory consumption varied widely among chromosomes for mm2-ivh-hpc and mm2-k133-hpc, whereas hifiasm-ont-r1 exhibited stable performance (Supplementary Section S4.2, Supplementary Tables S4–S6). In both mm2-ivh-hpc and mm2-k133-hpc, the overlap detection step consumed the dominant proportion of execution time. The total CPU time was 11.8 hours for mm2-ivh-hpc, 339.7 hours for mm2-k133-hpc, and 3.5 hours for hifiasm-ont-r1. The geometric mean of memory usage per chromosome (worst case in parentheses) was 6.4 GiB (14.0 GiB), 40.5 GiB (181 GiB), and 16.5 GiB (16.8 GiB) in the same order. The total size of the overlap detection outputs (PAF files) was 77.0 GiB and 1.08 TiB for mm2-ivh-hpc and mm2-k133-hpc, respectively (Supplementary Table S7).

## Discussion

The benchmark clearly demonstrated that interval hashing is effective for precise overlap detection in alpha satellite HORs. The substantial reductions in execution time, memory usage, and output file size for overlap detection compared to the control using standard minimizers (mm2-k133-hpc) also indirectly support that interval hashing reduces false-positive matches. These findings suggest that the intervals of k-mers within repeats are a critical feature for distinguishing alpha satellite HORs. We also noticed that the concept of k-mer intervals aligns with early studies of alpha satellites in which HORs are characterized by analyzing the lengths of DNA fragments after restriction enzyme digestion (Wu and Manuelidis 1980). This indicates that interval hashing essentially examines the same feature, renovating one of the most fundamental features of alpha satellite HORs using modern techniques.

Interval hashing resembles seed-and-extend alignment algorithms for optical mapping (Leung *et al*. 2017; Salmela *et al*. 2020) in that both use subsequences of interval values as seeds. Our contribution lies in demonstrating that subsequences of interval values are useful as signatures of repeats, which has been less explored in the field of optical mapping. The specific structure of interval hashing is also similar to RawHash, a hashing algorithm for matching on Nanopore raw signals, in that both construct hashes from coarsened numerical sequences (Firtina *et al*. 2023). In comparison with algorithms in the sequence space, interval hashing is similar to spaced seeds (Ma, Tromp and Li 2002) and strobemers (Sahlin 2021) in that these algorithms construct a single seed from multiple exact submatches separated by gaps. We found interval hashing and existing algorithms essentially differ in whether or not the submatches are synchronized with the underlying repeat structure. Interval hashing uses synchronized submatches, which effectively capture repeat contexts.

The weakness of interval hashing is its susceptibility to sensitivity degradation due to sequencing errors. This stems from the algorithm’s reliance on the preservation of all k-mers within *W* occurrences before and after a match. In our experiments, the low error rates of ONT reads appeared to mitigate the sensitivity issue.

Improvements to the algorithm will be necessary to apply interval hashing to data with higher error rates. In the field of optical mapping, algorithms for obtaining error-robust seeds on interval information have been explored (Salmela *et al*. 2020), which may help improve our algorithm.

Although our benchmark focused on alpha satellite HORs, interval hashing is likely to be compatible with unique genomic regions, segmental duplications, and isolated interspersed repeat elements. This compatibility stems from the property that singleton or isolated minimizers maintain the same values after applying interval hashing, thereby not affecting the behavior of the algorithms that use minimizers.

Moreover, interval hashing potentially enhances specificity in less structured repeat regions, such as various satellite repeats and clusters of interspersed repeats, similar to its effect on alpha satellite HORs. These properties could make interval hashing applicable to pipelines relying on all-versus-all overlaps of reads (i.e., without probing or filtering, such as whole-genome assembly and read error correction (Espinosa *et al*. 2024; Stanojevic *et al*. 2024)), potentially improving accuracy in various repeat contexts. Our implementation, mm2-ivh, facilitates evaluation of how interval hashing improves the performance in these pipelines on diverse data.

## Acknowledgments

The computational resources used in this study were provided by the Human Genome Center, the Institute of Medical Science, the University of Tokyo. We also thank Dr. Yoshihiko Suzuki for providing insightful comments on the algorithm and the manuscript.

## Author contributions

Hajime Suzuki: Conceptualization, Methodology, Software, Validation, Formal analysis, Investigation, Visualization, Writing – original draft. Masahiro Sugawa: Validation, Resources, Writing – review & editing. Yoshitaka Sakamoto: Validation, Writing – review & editing. Yuichi Shiraishi: Conceptualization, Resources, Writing – review & editing, Supervision, Funding acquisition.

