## Supplementary Notes for "mm2-ivh: simple and precise overlap detection in alpha satellite HORs with interval hashing"

### S1. mm2-ivh implementation details

#### S1.1. Building interval-hash-augmented minimizer index

To incorporate interval hashing into minimap2, we implemented a function `mm_idx_patch_ivh` that takes the minimap2 index data structure and transforms the stored minimizers into interval-hash-augmented minimizers. This function retrieves the minimizers and their occurrence position lists from the hash table in the index, and then calculates the interval hash values from the occurrence position lists. The original minimap2 index data structure stores the position lists sorted, allowing the interval vector to be constructed by subtracting adjacent elements. The resulting interval hash values are combined with the original minimizers using XOR to form augmented minimizers. The augmented minimizer list is then sorted and grouped by identical values, with each group inserted into a new hash table. In cases where augmented minimizer values collide, only the one with the fewest occurrences across the entire indexed sequence (target sequence) is retained. After processing all minimizers, the original minimizer hash table is replaced with the new augmented minimizer hash table.

The constructed augmented minimizer index is compatible with the rest of minimap2 when used within mm2-ivh. This means that the interval hashing can also be applied to various computations that minimap2 can perform, such as mapping to reference sequences or aligning whole genomes. The code for processing minimizers other than index construction uses the original minimap2 code, with only minor additional tweaks explained in the following sections S1.2 and S1.3.

#### S1.2. Filtering out locally frequent minimizers

Some repeat sequences may still retain excessively frequent minimizers even after applying interval hashing. We considered that locally high-frequency minimizers carry little information and that removing them strikes a good balance between practical sensitivity and computation time. To filter out locally high-frequency minimizers, mm2-ivh counts the occurrences of minimizers within a fixed-length window and removes those that exceed a certain threshold from the index. This filtering process is applied only when interval hashing is enabled. The default setting is to count occurrences within a 1000 bp window, removing minimizers that appear 24 times or more. Since the monomer of alpha satellite is 171 bp, this setting does not remove minimizers derived from monomers in typical alpha satellite HORs. However, for repeats composed of shorter elements such as beta satellite, human satellite, and tandem repeats, this filtering may reduce the sensitivity of overlap detection. The actual impact of this filtering on repeats other than alpha satellite is outside the scope of this paper and needs to be investigated in future research.

#### S1.3. Minimizer hit count compensation

Interval hashing requires that the positional relationships of minimizers encoded in the hash values are preserved for the augmented minimizers to match against the correct counterpart. This means that if a sequencing error disturbs the occurrence of at least one of the $W$ minimizers preceding or following a given minimizer, the augmented minimizer will fail to produce a true positive match.

This causes a drift in the estimation of the number of matching bases and the error rate (sequence divergence) on the overlap path, which are calculated from the hit counts of minimizers on the overlap chain in the original minimap2 algorithm. We implemented a compensation for the hit counts in mm2-ivh to mitigate this drift. For a given chain, it examines the augmented minimizers on the query sequence, specifically the $W$ augmented minimizers before and after the central augmented minimizer. If any of these have hits, it considers that the central minimizer also had a hit. This compensation is not perfect; it does not fully restore the hits obtained from the original minimizers and may introduce a small fraction of false positive hits. However, we confirmed using simulated reads that the approximate sequence divergence (`dv:f` tag) reported in the PAF records by mm2-ivh is closer to the value reported by the original minimap2 in unique regions when this compensation is applied, compared to when it is not (Supplementary Section S3.5).

### S2. Modification on minimap2 to support longer k-mers

In our main experiments, the interval hashing configuration in mm2-ivh-hpc requires a seed hit with 7 consecutive minimizers of length $k = 19$. This is derived from the interval hashing wing length of $W = 3$. For comparison with mm2-ivh-hpc, we used a standard minimizer (mm2-k133-hpc) with an equivalent k-mer length of $k = 7 \cdot 19 = 133$.

The original minimap2 implementation restricts the minimizer length ($k$) to a maximum of 28 due to the 56-bit field used to store minimizers. Since $k = 133$ exceeds this limit, we modified mm2-ivh, which is based on minimap2 2.28, to handle longer k-mers. The implementation is as follows: when a $k$ longer than 28 is specified, the k-mer is sliced from the lower bits in 49-bit chunks before calculating the minimizer hash value. The `hash64` function, which is also used in the original implementation to hash k-mers, is applied independently to each chunk. The resulting hash values are combined by left-shifting $i$-th chunk by $i$ bits and XORing them to obtain the final hash value. We implemented this hash function using the C23 `_BitInt` feature. Since C23 is a relatively new standard and is incompatible with older compilers, we did not include this implementation in the main mm2-ivh codebase. Instead, this implementation is placed in a derived branch for the experiment in this paper, identified by the `benchmark` tag. Additionally, since this implementation reduces the hash value to a shorter length than the original k-mer, it may introduce collisions with a low probability. However, we believe that this probability is sufficiently low for real datasets.

The index function of minimap2 excludes k-mers from the index if their span before homopolymer compression is 256 bases or more. If many k-mers are excluded with the $k = 133$ setting, it could unintentionally reduce the sensitivity of overlap detection. To confirm that this is not an issue for the HOR regions, we extracted all 256-base windows from the 500 kbp margined HOR regions of HG002 (see Evaluation section) in a sliding window manner and applied homopolymer compression to each window. We counted the fragments that were 133 bases or shorter after compression, finding that their proportion was only 0.0068%. Additionally, the longest continuous segment of fragments that were 133 bases or shorter was 345 windows long, indicating that the number of minimizers excluded due to homopolymer compression in the HOR regions of HG002 when using $k = 133$ is negligible.

### S3. Preliminary Experiments

Prior to the main experiments presented in the paper, we conducted preliminary experiments using simulated reads to quantitatively evaluate the sensitivity and precision of interval hashing in local diploid assembly. We selected HOR regions from two chromosomes as assembly targets: chromosome 1, which is relatively easy to assemble, and chromosome 7, which is more challenging.

#### S3.1. Methods

##### *S3.1.1. Preparation of template sequences for read simulation*

Simulated reads were generated from the alpha satellite HOR regions of HG002 v1.1 to match our main experiments. The sequence of HG002 v1.1 was downloaded from https://s3-us-west-2.amazonaws.com/human-pangenomics/T2T/HG002/assemblies/hg002v1.1.fasta.gz. The alpha satellite HOR regions of HG002 v1.1 were defined by extracting records starting with `active_hor` in the name field (4th column) from the annotation file downloaded from https://s3-us-west-2.amazonaws.com/human-pangenomics/T2T/HG002/assemblies/annotation/centromere/hg002v1.1_v2.0/hg002v1.1.cenSatv2.0.bed, and adding a 100 kbp margin to those regions. The extracted sequences from these regions are referred to as the margined HOR sequences of chromosome 1 and 7, respectively.

Additionally, to evaluate the effect of minimizer hit count compensation, we also used unique regions without complex repeats. We selected the region 50,000,000-55,000,000 on both chr1_MATERNAL and chr1_PATERNAL of HG002 v1.1 for this purpose. The extracted sequence from this region is referred to as the unique sequence of chromosome 1.

#### *S3.1.2. Overlap detection configurations*

We defined two overlap detection configurations to be used throughout the subsequent preliminary experiments. Both configurations are based on the `-x ava-ont` preset of minimap2, with homopolymer compression (HPC) enabled and the minimizer window size $w$ set to 15.

1. A configuration using mm2-ivh with interval hashing enabled. In this configuration, the minimizer length $k$ was fixed at 19, and the wing length $W$ was varied. The maximum interval for interval hashing was set to 20 kbp, and the options related to filtering locally frequent minimizers were set to the defaults described in Supplementary Section S1.2. We refer to this configuration as “mm2-ivh-hpc” in the subsequent preliminary experiments.
2. A configuration using the modified minimap2 that supports longer k-mers as described in Supplementary Section S2. In this configuration, the minimizer length was varied, and the wing length $W$ was fixed at $W = 0$ (disabling interval hashing). The filter for locally frequent minimizers was disabled in this configuration. We refer to this configuration as “mm2-k-hpc”.

#### *S3.1.3. Read simulation and overlap detection*

We conducted two experiments to investigate the quantitative characteristics of interval hashing. In these experiments, we varied two factors that are considered to most significantly impact the quality of the overlaps reported: the wing length $W$ of interval hashing and the quality value (QV) of the reads. The two experiments are defined as follows:

1. The mean quality value (QV) of the reads is fixed, and the wing length $W$ of interval hashing is varied. This experiment is referred to as “varW.”
2. The QV of the reads is varied, and the wing length $W$ of interval hashing is fixed. This experiment is referred to as “varQV.”

We defined two simulation profiles to generate reads for the two experiments, representing typical High-Fidelity (HiFi) reads and Oxford Nanopore Technologies (ONT) ultra-long (UL) reads. These profiles specify three aspects: read length distribution, error model, and quality score model. The error model and quality score model are those defined by Badread (https://github.com/rrwick/Badread; commit f38ef6f), which we used for read generation in the preliminary experiments. The details of the two profiles are as follows:

1. A profile that represents the length distribution of PacBio HiFi reads, with a mean length of 15 kbp, standard deviation of 4 kbp, and the error model and quality score model pacbio2021. This profile is referred to as “hifi.”
2. A profile that represents the length distribution of ONT UL reads, with a mean length of 25 kbp, standard deviation of 50 kbp, and the error model and quality score model nanopore2023. This profile is referred to as “ul.”

For the varW experiment, we generated the following three simulated samples from the margined HOR sequences of chromosomes 1 and 7, respectively:

1. A sample generated using the hifi profile with a mean QV of 27 (corresponding to a mean identity of 99.80%). Maximum identity and identity standard deviation (SD), which are required by Badread, were set to 99.995 and 0.100, respectively. This sample is referred to as “hifi-hor-QV27” for both chromosomes 1 and 7.
2. A sample generated using the ul profile with a mean QV of 17 (identity of 98.00%). Maximum identity and identity SD were set to 98.200 and 0.100, respectively. This sample is referred to as “ul-hor-QV17.”

3. A sample generated using the ul profile with a mean QV of 25 (identity of 99.68%). Maximum identity and identity SD were set to 99.720 and 0.016, respectively. This sample is referred to as "ul-hor-QV25."

The target coverage for all samples was set to 15x. We also disabled options that mimic real-world mechanisms of errors, i.e. junk reads, random reads, chimeras, and glitches. On the three generated samples, we performed overlap detection using mm2-ivh-hpc varying the wing length $W$ from 0 to 4. As a comparison, we also tested mm2-k-hpc with the k-mer length set to $k = 19 \cdot (2W + 1)$ to match the number of matching bases of mm2-ivh-hpc for $W = 0...4$. Specifically, $k = 19, 57, 95, 133$, and 171 correspond to $W = 0, 1, 2, 3$, and 4, respectively.

For the varQV experiment, we generated a set of 11 samples from the margined HOR sequences of chromosomes 1 and 7 using the ul profile. The 11 samples were generated with different mean QVs, ranging from 10 (corresponding to an identity of 90%) to 34 (identity of 99.96%). The specific values, along with maximum identity and identity SD, are provided in Supplementary Table S1. The sample set is referred to as ul-hor-varQV for both chromosomes 1 and 7. In these two sample sets, we performed overlap detection using only mm2-ivh-hpc, which uses the minimizer parameters $(k, w) = (19, 15)$. The wing length $W$ was fixed at 3. We did not test mm2-k-hpc in this experiment due to its significantly longer computation time.

As a derivative of the varQV experiment, we also conducted an experiment to evaluate the effectiveness of the minimizer hit count compensation described in Supplementary Section S1.3. For this experiment, we generated a set of 11 samples from the unique sequence of chromosome 1 in the same manner as ul-hor-varQV. This sample set is referred to as ul-uniq-varQV. Along with ul-hor-varQV, we performed overlap detection under the following three configurations:

1. The varQV experiment using ul-hor-varQV as described above.
2. The varQV experiment using ul-hor-varQV with hit count compensation disabled. We used the version of mm2-ivh tagged `preliminary-experiments`, which reports the approximate sequence divergence without hit count compensation under a separate tag `rd:f`.
3. A configuration using ul-uniq-varQV and detecting overlaps with mm2-k-hpc with $k = 19$. This is equivalent to running the original minimap2 with the `-x ava-ont` preset. This configuration represents the case where all hits for $k = 19$ are obtained for reads with the same QV as the previous two configurations.

For each of the 11 samples in the results, we calculated the mean and variance of the approximate sequence divergence, which is a metric directly computed from the minimizer hit counts. The mean and variance were calculated from all overlaps without any filtering.

#### *S3.1.4. Modifications to the paftools ov-eval subcommand*

For evaluation in the two experiments, varW and varQV, we measured sensitivity and precision. We used a modified version of the ov-eval subcommand from paftools, which is distributed with minimap2. The modified paftools can be found in the mm2-ivh repository at the commit tagged `preliminary-experiments`. The specific modifications we made are as follows:

1. Verifying if the reported overlaps are sufficiently long and on the correct diagonal.
2. Reporting precision in addition to sensitivity.

In the first modification, we considered an overlap to be correct only if it was at least 80% of the length of the true overlap. Overlaps are often fragmented in repeat regions, and fragmented overlaps can lead to assembly breaks. By considering short overlaps as incorrect, we prevented the overestimation of outputs that contain many fragmented overlaps. In addition, we checked whether the overlaps were reported on the correct diagonal. Long false overlaps are often detected parallel to the true overlaps in repeat regions. If all of these were considered correct, sensitivity could be overestimated. We considered an overlap to be correct only if the diagonal deviation of endpoints of the reported overlap from the corresponding endpoints of the correct overlap was within 2 kbp.

In the second modification, we added precision calculation in addition to sensitivity. We calculated precision for two types of overlap sets: 1) the set of all reported overlaps of 1.6 kbp or longer, and 2) its subset, which retains only the longest overlap among multiple overlaps reported for the same read pair. The former represents general precision and the latter is intended as a metric to measure compatibility with string graph assemblers. The threshold of 1.6 kbp for the former was determined to be consistent with both the original paftools ov-eval implementation and our first modification. The original paftools ov-eval infers the true overlap set from intersections of read mappings that are 2 kbp or longer on the reference. Our first modification considers only overlaps that are at least 0.8 times the length of the true overlap as correct. To define an overlap set for precision measurement that is consistent with both, we extracted the subset of reported overlaps that are 1.6 kbp ( $= 2 \cdot 0.8$ kbp) or longer. The latter subset was created by removing non-longest overlaps from the former set when multiple overlaps were reported between the same read pair. The string graph algorithm simplifies the overlap graph by retaining longer overlaps when two overlaps are conflicting or redundant. That is, longer overlaps are more likely to mislead the final assembly if they are incorrect. We considered the correctness, especially precision, of the subset retaining only the longest overlaps is a concise metric to assess whether the string graph algorithm can produce a correct assembly. Hereafter, we refer to the former simply as “precision” and the latter as “precision in the longest overlap subset” for distinction.

##### *S3.1.5. Plotting sensitivity-precision curves*

Using the results of the varW experiment, we plotted sensitivity-precision curves by continuously varying the threshold for filtering the reported set of overlaps. As the filtering criterion, we used the fraction of matching bases to the total overlap length, i.e., the match rate. Specifically, we obtained the value by dividing the match length field (10th column) by the block length field (11th column). We considered this to be a sufficiently concise and universal criterion, as it is calculated solely from fields defined in the PAF format specification and does not depend on optional tags. The filtering was performed to retain only overlaps with a match rate above the threshold. The match rate threshold was varied from 0.0 to 1.0 in increments of 0.005, and paftools ov-eval was executed on each resulting overlap subset to obtain sensitivity and precision. Here, precision refers to the former of the two types of precision mentioned earlier, measured on the set of all overlaps of 1.6 kbp or longer.

#### S3.2. Results

The sensitivity in the varW setting is shown in Supplementary Figure S1. In the hifi-hor-QV27 profile, enabling interval hashing, i.e., $W \geq 1$, resulted in a decrease in sensitivity. In the ul-hor-QV17 profile, a decrease in sensitivity was observed for $W \geq 3$. In the ul-hor-QV25 profile, the decrease in sensitivity was minimal compared to the other two profiles. In all profiles, chromosome 7 exhibited a greater decrease in sensitivity compared to chromosome 1.

The sensitivity-precision curves for mm2-ivh-hpc with $W = 3$ and mm2-k-hpc with $k = 133$, extracted from the varW experiment, are shown in Supplementary Figure S2. mm2-ivh-hpc with $W = 3$ improved precision compared to mm2-k-hpc with $k = 133$, albeit with a slight to moderate decrease in sensitivity except for centromere 7 of ul-hor-QV17. In the ul-hor-QV25 sample for chromosome 1, the decrease in sensitivity was negligible, resulting in a purely improved precision. The precision in the longest overlap subset is shown in Supplementary Table S2. mm2-ivh-hpc achieved the precision exceeding 99% for chromosome 1, while it remained around 70% for chromosome 7. mm2-k-hpc achieved approximately 30% and 20% for chromosomes 1 and 7, respectively.

The changes in sensitivity in the varQV setting are shown in Supplementary Figure S3. The interval hashing with parameters set to $(k, w, W) = (19, 15, 3)$ exhibited sufficiently high sensitivity when the mean QV of the reads was 20 or higher. In this experiment as well, chromosome 1 consistently exhibited higher sensitivity than chromosome 7.

Supplementary Figure S4 shows the distribution of approximate sequence divergence in the read sets of ul-hor-varQV and ul-uniq-varQV. The minimizer hit count compensation brought the mean value of approximate sequence divergence in ul-hor-varQV closer to that of ul-uniq-varQV. Comparing chromosome

1 and chromosome 7, chromosome 1 exhibited a smaller standard deviation in approximate sequence divergence and a smaller deviation between ul-hor-varQV and ul-uniq-varQV.

#### S3.3. Discussion

In the varW experiment, hifi-hor-QV27 exhibited a decrease in sensitivity even with the shortest wing length interval hashing, $W = 1$. The results of ul-hor-QV17 and ul-hor-QV25 showed relatively high sensitivity despite having lower identity than the hifi-hor-QV27 sample. These results indicate that for interval hashing to achieve sufficient sensitivity, the reads need to be sufficiently long. Comparing ul-hor-QV25 and ul-hor-QV17, which have the same read length distribution, a decrease in sensitivity was observed only in ul-hor-QV17 for larger $W$ (approximately $W \geq 3$), suggesting a trade-off between sensitivity and $W$. The varQV experiment demonstrated a specific manifestation of this trade-off at $W = 3$, indicating that a QV of approximately 20 (corresponding to 99% identity) or higher is required.

The sensitivity-precision plots from the varW experiment suggest that interval hashing significantly improves precision at the cost of a slight decrease in sensitivity. The precision in the longest overlap subset indicates that the improvement in precision is achieved in a manner compatible with the string graph algorithm. Notably, the precision exceeding 99% for chromosome 1 may suggest that the overlaps obtained using interval hashing allow for accurate assembly without any additional post-processing of the overlaps.

The distribution of approximate sequence divergence in the varQV experiment suggests that minimizer hit count compensation can partially compensate for hits lost due to interval hashing in HOR regions. However, there remains a deviation from the values reported in unique regions, which can be considered as the values when all minimizers correctly hit, indicating that the compensation is not complete.

Based on the above results, we concluded that the parameters $(k, w, W) = (19, 15, 3)$ can achieve practical sensitivity and precision with UL reads of approximately QV = 20. We adopted this value as the default for the `-x ivh-ava-ont-ul` preset.

### S4. Main experiments

In this section, we describe the details of the methods of the experiments presented in the main text.

#### S4.1. Input preparation

We used the CHM13 v2.0 reference genome published by the Telomere-to-Telomere consortium for read binning, downloaded from https://s3-us-west-2.amazonaws.com/human-pangenomics/T2T/CHM13/assemblies/analysis_set/chm13v2.0.fa.gz. The corresponding repeat annotation file for CHM13 v2.0 was taken from https://s3-us-west-2.amazonaws.com/human-pangenomics/T2T/CHM13/assemblies/annotation/chm13v2.0_censat_v2.1.bed. We extracted records starting with `hor` in the name (4th column), merged those that were within 5 Mbp of each other, and added a 500 kbp margin. The resulting regions are referred to as "HOR regions with 500 kbp margins" in the main text.

The ultra-long (UL) reads used in the benchmark is the high-accuracy UL sequencing data provided by Oxford Nanopore Technologies on the EPI2ME blog (https://epi2me.nanoporetech.com/gm24385_ncm23_preview/). The data was downloaded from the URL s3://ont-open-data/gm24385_2023.12/all_pass.vhg002v1.bam. As a preprocessing step, we extracted the sequence information from the BAM file and converted it to FASTQ format. This resulted in a FASTQ file of 5,552,000 reads with a total base count of 126.9 Gbp (mean length 22.9 kbp) and a mean QV of 23.73. We then filtered the FASTQ file using Filtlong (commit 7c654f1), keeping only reads with a mean QV of 10.0 or higher and a length of at least 50 kbp. The filtered FASTQ file contained 760,699 reads with a total base count of 89.0 Gbp (mean length 116.9 kbp) and a mean QV of 24.64.

The filtered FASTQ file was mapped to the aforementioned CHM13 reference using minimap2 2.28 with the `-x map-ont` preset. Secondary mapping records (those with the `tp:A` tag not set to `P`) were removed. Reads that had an overlap of at least one base with the 500 kbp margined HOR regions were assigned to those regions (chromosomes). If a single read split and mapped to multiple different chromosomes as primary mappings, that read was assigned in full length to all the chromosomes without trimming. The number of reads and total base count per chromosome (assembly bin) are shown in Supplementary Table S3.

#### S4.2. Assembly

We created a script `run_local_assembly.sh` to perform read binning using the reference sequences and execute local assembly for each bin. The binning step in this script corresponds to mapping to the reference and the subsequent assignment of reads explained in the previous section. The script runs in either mode to execute programs that have input/output file formats and options compatible with minimap2 and miniasm (`-m minimap2`), or to execute Hifiasm (`-m hifiasm`). In the minimap2 mode, the minimap2-compatible program is configured not to compute the CIGAR tag. The script is located in the `scripts` directory of the https://github.com/ocxtal/mm2-ivh repository, and the version used in the benchmark is tagged with `benchmark`. An archive of this version is also available at 10.5281/zenodo.17143522. The options provided to the script were as follows:

```bash
# mm2-ivh
bash run_local_assembly.sh \
    -m minimap2 \
    -b mm2-ivh \
    -e "-xivh-ava-ont-ul -H -k19 -w15 --wing=3 --dual=no" \
    -E "-h5000 -c2" \
    -d run_mm2-ivh-hpc \
    -t 32 \
    -r /data/chm13/v2.0/chm13v2.0.fa.gz \
    -a /data/chm13/v2.0/chm13v2.0_censat_v2.1.bed \
    -p all.flt.m50k.90.chm13.v2.0.xmapont.paf \
    -q /data/hg002_q26/all.flt.m50k.90.fq.gz

# mm2-k133-hpc
bash run_local_assembly.sh \
    -m minimap2 \
    -b mm2-ivh \
    -e "-xivh-ava-ont-ul -H -k133 -w15 --wing=0 --dual=no" \
    -E "-h5000 -c2" \
    -d run_mm2-k133-hpc \
    -t 32 \
    -r /data/chm13/v2.0/chm13v2.0.fa.gz \
    -a /data/chm13/v2.0/chm13v2.0_censat_v2.1.bed \
    -p all.flt.m50k.90.chm13.v2.0.xmapont.paf \
    -q /data/hg002_q26/all.flt.m50k.90.fq.gz

# hifiasm-ont-r1
bash run_local_assembly.sh \
    -m hifiasm \
    -b hifiasm \
    -e "--ont -r1" \
    -d run_hifiasm-ont-r1 \
    -t 32 \
    -r /data/chm13/v2.0/chm13v2.0.fa.gz \
    -a /data/chm13/v2.0/chm13v2.0_censat_v2.1.bed \
    -p all.flt.m50k.90.chm13.v2.0.xmapont.paf \
    -q /data/hg002_q26/all.flt.m50k.90.fq.gz
```

The CPU time and maximum memory usage consumed during the assembly step are shown in Supplementary Tables S4–S6. These values were obtained by parsing the logs output by the tools. The total size of the intermediate output files (pairwise alignment format (PAF) files) is shown in Supplementary Table S7.

#### S4.3. Evaluation

For evaluating the assembly results, we plotted dotplots of the assembled unitigs against the alpha satellite HOR regions from HG002 v1.1, which is published by the Telomere-to-Telomere consortium. The HG002 v1.1 assembly was downloaded from https://s3-us-west-2.amazonaws.com/human-pangenomics/T2T/HG002/assemblies/hg002v1.1.fasta.gz. The alpha satellite HOR regions were extracted from the annotation file by selecting records starting with `hor` or `active_hor` in the name (4th column), merging those that were within 5 Mbp of each other, and adding a 500 kbp margin. The annotation file we used was downloaded from https://s3-us-west-2.amazonaws.com/human-pangenomics/T2T/HG002/assemblies/annotation/centromere/hg002v1.1_v2.0/hg002v1.1.cenSatv2.0.bed. The dotplots were created using our own implementation called tenten (https://github.com/ocxtal/tenten), utilizing only minimizers with a count of 5 or fewer across the entire 500 kbp margined HOR regions. Generated dotplots are included in the `main_experiment` directory of the Supplementary Files.

We visually inspected the dotplots to assess the consistency between the assembly results and HG002 v1.1. The results of this evaluation are presented in Supplementary Table S8. The evaluation focused on whether a single unitig covered the active HOR region (records with `active_hor` in the annotation) and whether it contained any misjoins (confusion between haplotypes or connections at different positions). The presence or correctness of edges connecting unitigs and misassemblies in regions other than the active HOR regions were ignored. The reason we focused only on the active HOR regions and did not consider the other HOR regions (records with `hor`) is that some chromosomes have fragmented HOR regions. This makes it difficult to determine whether misassemblies in those regions are due to the characteristics of the assembler or other reasons, such as the inability to recover reads corresponding to the gaps between CHM13 and HG002 HORs. In contrast, the active HOR regions are single or sufficiently large even if split into multiple parts, making it easier to assess the characteristics of overlap detection in alpha satellite HOR regions.

#### S4.4. Further inspection of misjoin in the Hifiasm output

We identified a misjoin (haplotype switch) in the Hifiasm output (hifiasm-ont-r1) between chromosome 21 maternal and paternal. The corresponding coordinates on HG002 v1.1 were approximately 13.15 Mbp for chromosome 21 maternal and 9.55 Mbp for chromosome 21 paternal. We extracted two reads each from chromosome 21 maternal and paternal that were mapped across this boundary using minimap2 2.28 with the `-x map-ont` preset on the UL read data used for assembly. The correspondence between the names mat1/2 and pat1/2 in Figure 1(e) and the original read IDs, along with their mapped coordinates, is shown in Supplementary Table S9.

The dotplots in Figure 1(e) for “Interval-hash-augmented minimizers” and “Standard minimizers” were generated using the minimizer parameters in mm2-ivh-hpc and hifiasm-ont-r1, respectively. Specifically, the values were $(k, w, W) = (19, 15, 3)$ for mm2-ivh-hpc and $(k, w) = (51, 51)$ for hifiasm-ont-r1, with homopolymer compression enabled in both cases. The tool tenten was used for plotting, similar to the evaluation step, with a resolution of 2000 bp/pixel. Only minimizers with a count of 10 million or fewer were used in the plot. Since the total length of the query and target sequences does not exceed 10 Mbp, this effectively corresponds to plotting all minimizers. Note that the minimizers plotted are not the same as those used in the actual mm2-ivh and Hifiasm programs to compute overlaps, which have a filtering step to exclude frequently occurring minimizers.

To gain a more comprehensive understanding of how the seed matches are distributed, we present a complete version of Figure 1(e) in Supplementary Figure S5, which includes plots using minimizers with the parameters $(k, w) = (133, 15)$ corresponding to the mm2-k133-hpc setting in addition to those used in Figure 1(e). In this figure, all combinations of the four reads are shown. For each setting, the left panel shows all minimizers without filtering out frequently occurring ones, while the middle and right panels show minimizers with counts of 200 or fewer and 10 or fewer across the entire four reads, respectively. The resolution is the same as in Figure 1(e), set to 2000 bp/pixel. From the comparison of panel (a) with panel (b) or (c) in Supplementary Figure S5, it is indicated that interval hashing achieves seed matches with higher specificity compared to standard minimizers and that this specificity is maintained even when high-frequency minimizers are removed.

#### S4.5. Results of local assembly using sample-matched reference

In the experimental setup described in the main text, we used the CHM13 v2.0 reference sequence for binning the HG002 reads. This was intended to reflect the reality of local assembly problems, where a reference assembled from the same sample is typically not available. In this section, we present the results using a reference assembled from the same sample as the assembly target (referred to as “sample-matched reference”), i.e., a setting without bias from binning.

The methods for the experiment are the same as those described in Supplementary Sections S4.1–S4.3, except that HG002 v1.1 was used for binning. Since HG002 v1.1 contains both paternal and maternal sequences for autosomes, reads assigned to each were grouped into the same autosomal bin during binning. Chromosome X and chromosome Y were assigned to separate bins, as in the main experiment.

The number of reads collected from binning is shown in Supplementary Table S10. The differences from the CHM13 v2.0 reference were not significant, except for the acrocentric chromosomes (13, 14, 15, 21, and 22) and chromosomes 1, 3, and 4, where changes of 20% to 40% were observed. Computation time, memory usage, and the size of the PAF files for overlap detection output are shown in Supplementary Tables S11–S14. These showed consistent trends with those observed in the CHM13 v2.0 reference. In chromosomes where the number of reads changed, computation time, memory usage, and PAF file size all changed within a range consistent with the change in the number of reads, which was roughly proportional to the square of the change rate in the number of reads.

The manual inspection results of the assembly are shown in Supplementary Table S15. The dotplots used for inspection are included in the `sample_matched_reference` directory of the Supplementary Files. mm2-ivh-hpc recovered the full length of the active HOR regions in 24 haplotypes, which was three haplotypes fewer than in the CHM13 v2.0 reference. Three types of classification changes were observed: those that were missing in CHM13 v2.0 but became complete in HG002 v1.1 (chromosome 13 paternal), those that were complete but became near-complete (chromosome 7 maternal, chromosome 11 maternal, and chromosome 17 maternal), and those that were complete but became split (chromosome 4 maternal). hifiasm-ont-r1 recovered active HOR regions in 14 haplotypes, which is two fewer than the CHM13 v2.0 reference. There were five haplotypes (chromosome 13 paternal, chromosome 14 maternal, chromosome 15 maternal, chromosome 21 paternal/maternal) that changed from incomplete (collapse or missing) in the CHM13 v2.0 reference to complete in the HG002 v1.1 reference, and seven haplotypes (chromosome 1 maternal, chromosome 8 maternal, chromosome 13 maternal, chromosome 15 paternal, chromosome 18 paternal, and chromosome 20 paternal/maternal) that changed in the opposite direction. mm2-k133-hpc recovered the full length of the active HOR region only for chromosome Y, which is the same as in the CHM13 v2.0 reference. In the other haplotypes where mm2-k133-hpc could not recover the full length of the active HOR region, transitions among split, collapse, and missing were observed.

The reason mm2-ivh-hpc and hifiasm-ont-r1 recovered the full length of chromosome 13 paternal is likely that the sample-matched reference reduced misses in read binning for chromosome 13. On the other hand, it is unclear why mm2-ivh-hpc could not recover the full length in some of the other haplotypes. The changes were minor (either the contig became shorter or split), and we believe these are within the range of fluctuations caused by changes in the collected reads. The changes observed in hifiasm-ont-r1 were larger than those observed in mm2-ivh-hpc, with some haplotypes transitioning between complete and missing or collapse. The cause is also unclear, but it might be due to Hifiasm excessively correcting the reads. We suspected that the error correction step of Hifiasm might lead to one haplotype being completely overwritten by the other. This interpretation could be consistent with the observation that hifiasm-ont-r1 recovered the full length of the active HOR regions only in one haplotype for chromosomes 13 and 15, on opposite sides between CHM13 v2.0 and HG002 v1.1.

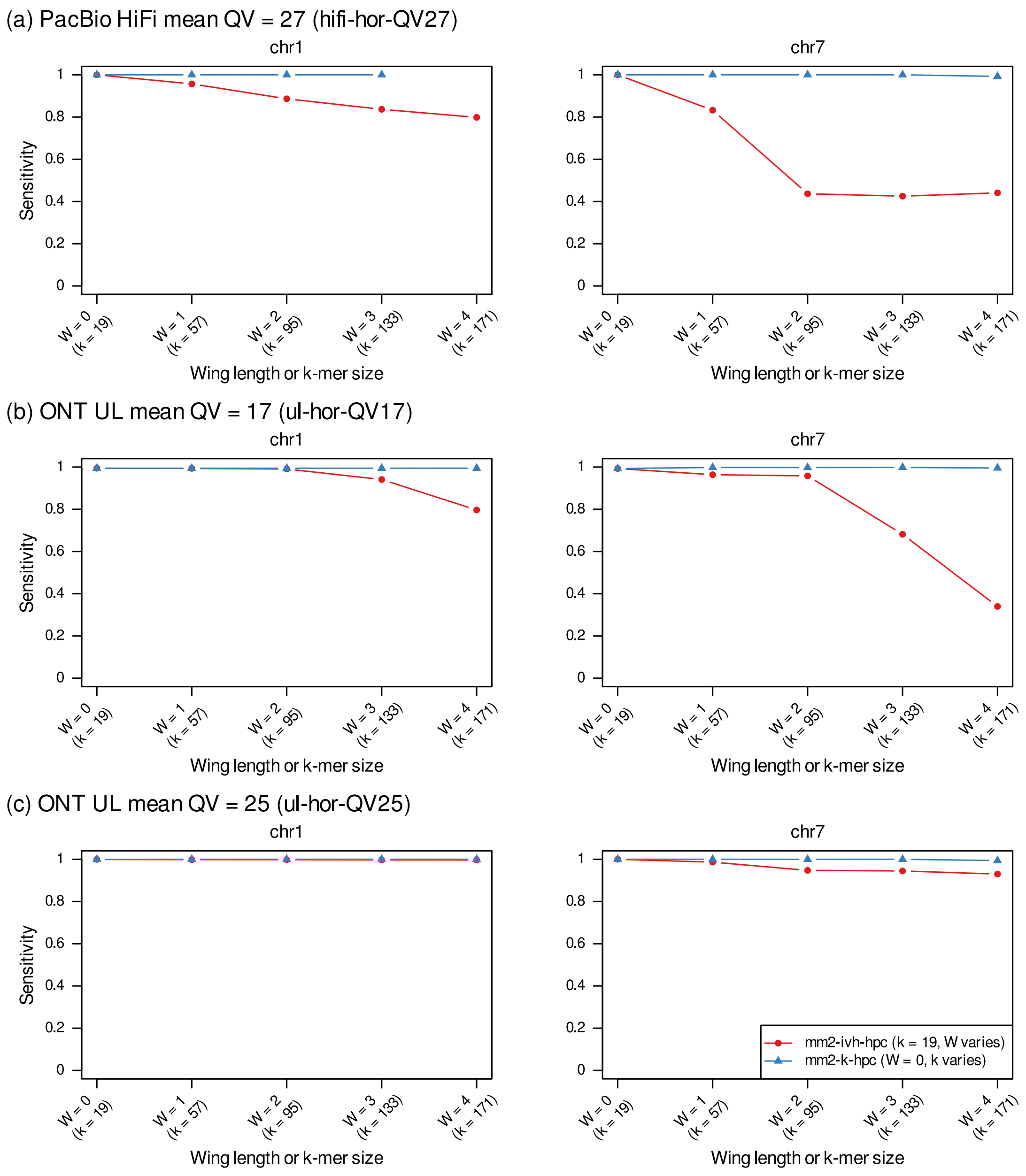


**Supplementary Figure S1. (a)** Sensitivity of overlap detection using mm2-ivh-hpc and mm2-k-hpc in the hifi-hor-QV27 sample, shown for chromosome 1 (left panel) and chromosome 7 (right panel). The x-axis labels indicate *W* for mm2-ivh-hpc outside the parentheses and *k* for mm2-k-hpc inside the parentheses. The point for $k$ = 171 of mm2-k-hpc in chromosome 1 is missing because the computation could not be completed due to insufficient disk space in the environment used. **(b)** Results for the ul-hor-QV17 sample in the same format. **(c)** Results for the ul-hor-QV25 sample in the same format.

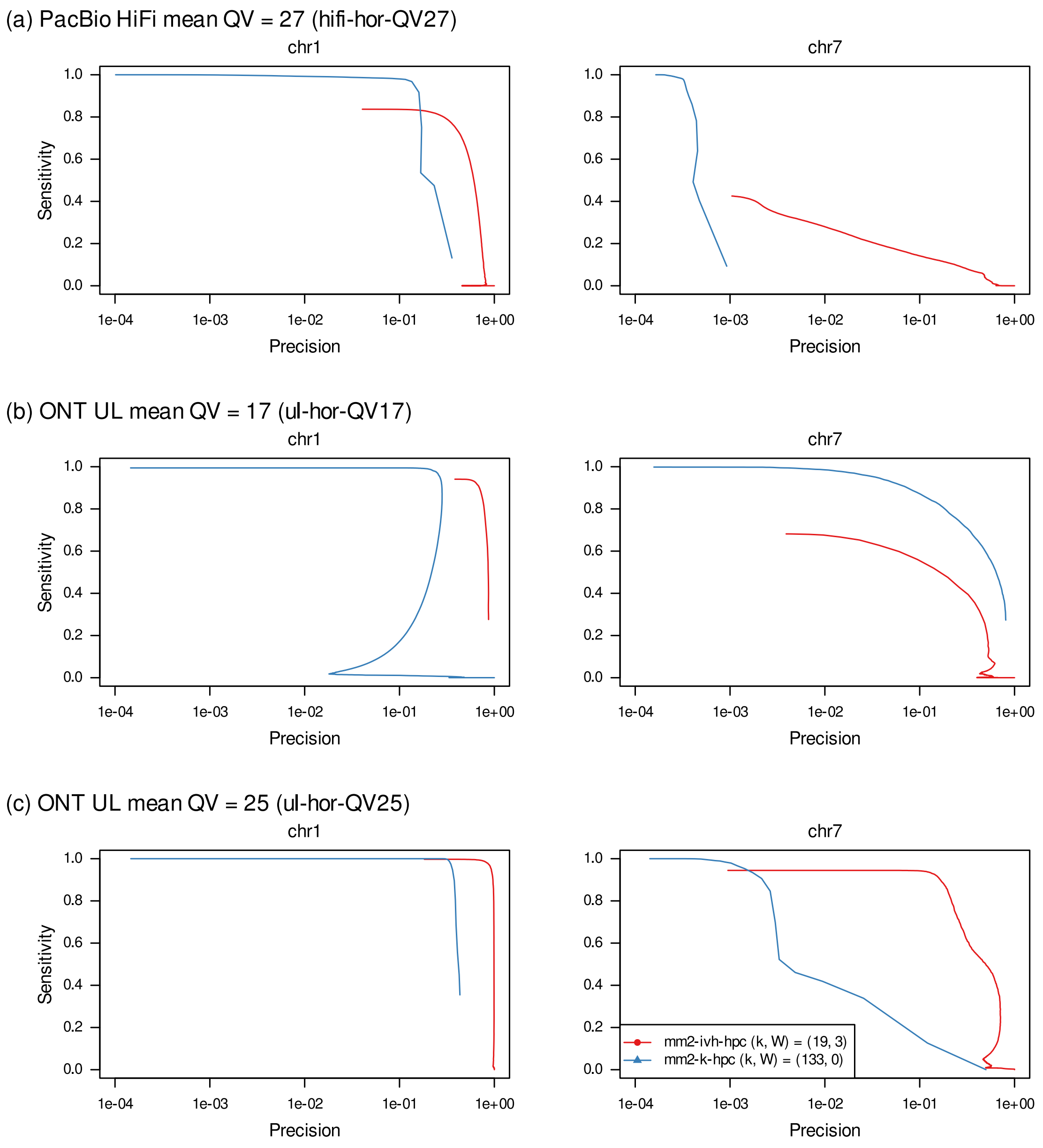


**Supplementary Figure S2. (a)** Sensitivity-precision curves for overlaps reported by mm2-ivh-hpc with $W = 3$ and mm2-k-hpc with $k = 133$ in the hifi-hor-QV27 sample. Results for chromosome 1 are shown in the left panel and for chromosome 7 in the right panel. **(b)** Results for the ul-hor-QV17 sample in the same format. **(c)** Results for the ul-hor-QV25 sample in the same format.

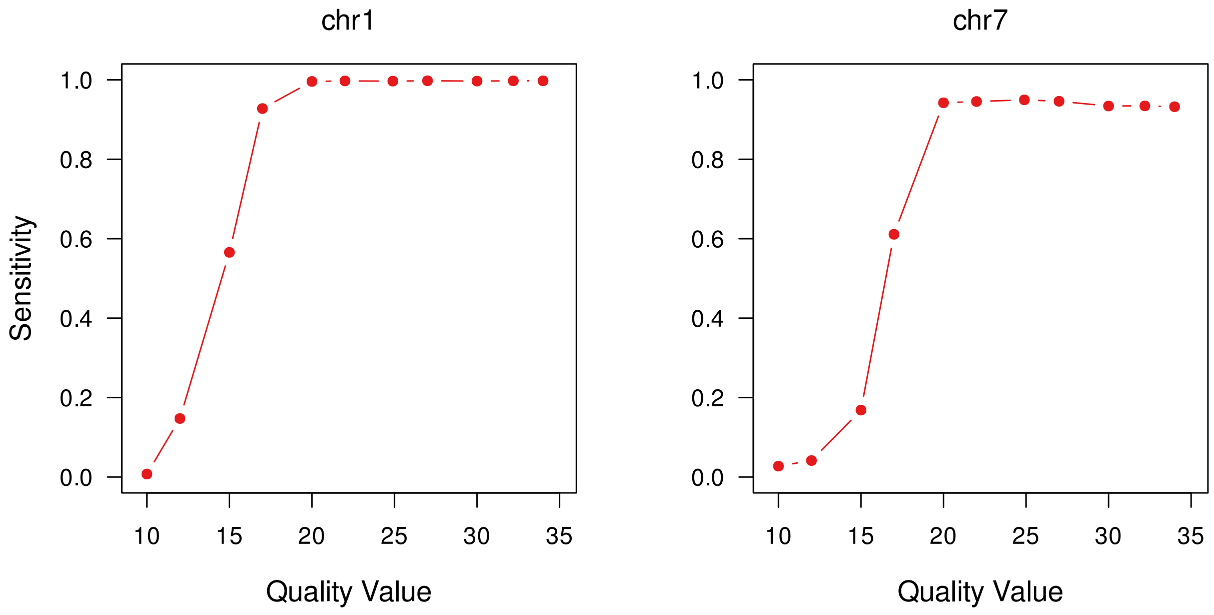


**Supplementary Figure S3.** Sensitivity of overlap detection using mm2-ivh-hpc with $W = 3$ in the ul-hor-QV25 sample set. Results for chromosome 1 are shown in the left panel and for chromosome 7 in the right panel. Mean QV values of 15, 20, and 25 correspond to mean identities of 96.83%, 99.00%, and 99.68%, respectively.

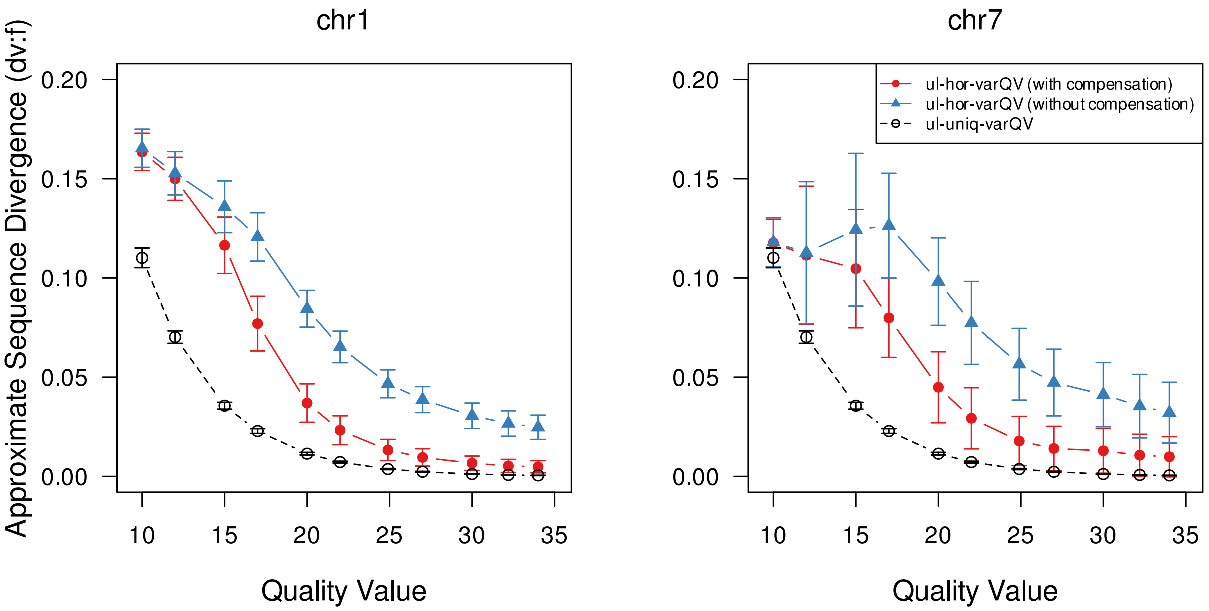


**Supplementary Figure S4.** Approximate sequence divergence (dv:f) reported by mm2-ivh and minimap2. ul-hor-varQV (with compensation) represents values when minimizer hit count compensation is enabled in mm2-ivh-hpc with $W = 3$ in the ul-hor-QV25 sample set. ul-hor-varQV (without compensation) represents values when minimizer hit count compensation is disabled in the same setting. ul-uniq-varQV shows values reported in the ul-uniq-QV25 sample set instead of ul-hor-QV25, with mm2-k-hpc with $k = 19$ (corresponding to the minimap2 `-x ava-ont` preset). Results for chromosome 1 are shown in the left panel and for chromosome 7 in the right panel.

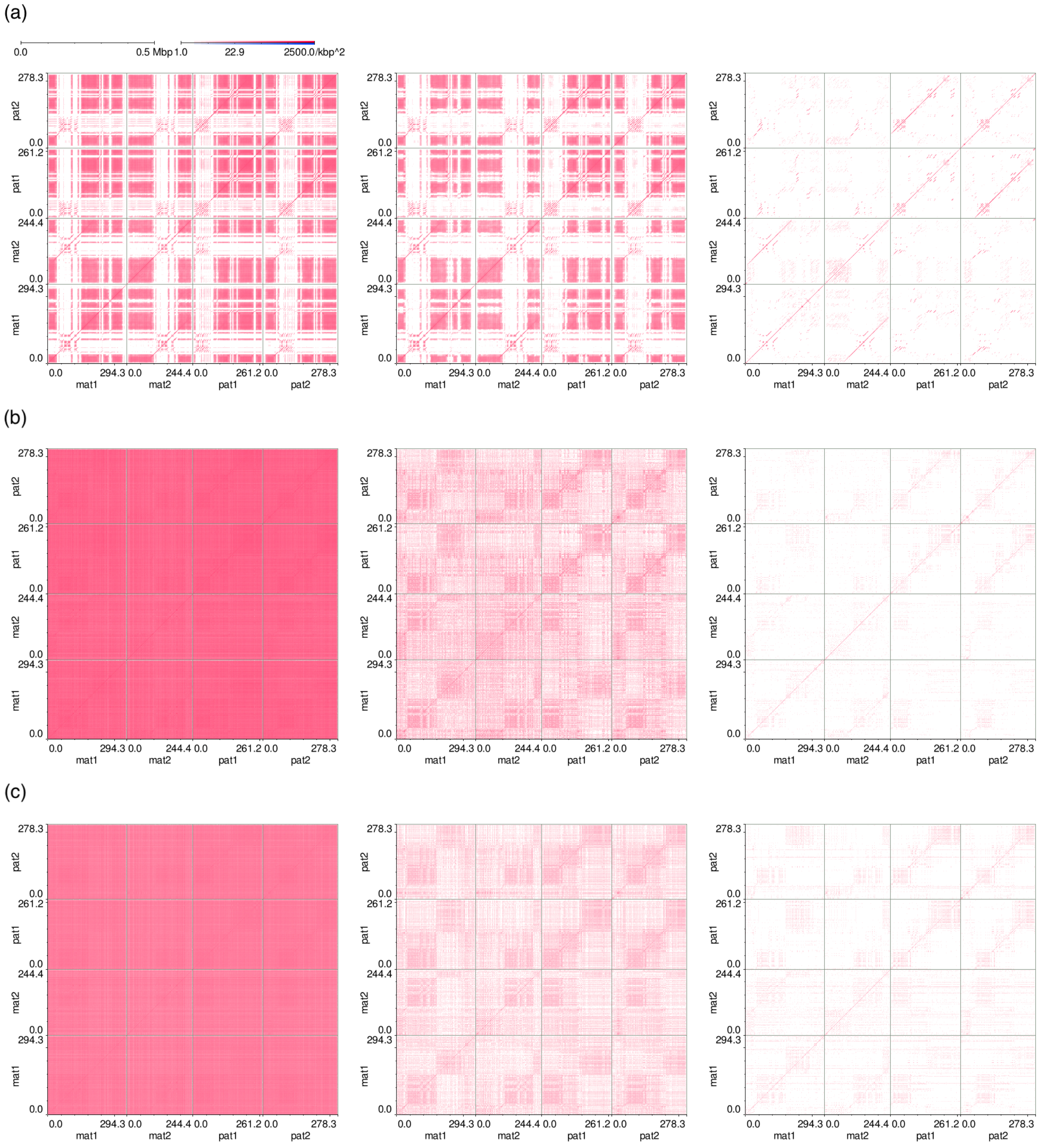


**Supplementary Figure S5.** **(a)** Interval-hash-augmented minimizers with $(k, w, W) = (19, 15, 3)$. **(b)** Standard minimizers with $(k, w) = (133, 15)$. **(c)** Standard minimizers with $(k, w) = (51, 51)$. Left panels show all minimizers, middle panels show minimizers with counts of 200 or fewer, and right panels show minimizers with counts of 10 or fewer across all four reads.

**Supplementary Table S1.** Quality values and corresponding mean identities used to generate the ul-hor-varQV sample set. Maximum identity and identity standard deviation (SD) given to Badread are also shown.

| Quality value (QV) | Mean identity (%) | Maximum identity (%) | Identity SD (%) |
|---|---|---|---|
| 10.0 | 90.00 | 91.00 | 0.500 |
| 12.0 | 93.69 | 94.32 | 0.315 |
| 15.0 | 96.84 | 97.15 | 0.158 |
| 17.0 | 98.00 | 98.20 | 0.100 |
| 20.0 | 99.00 | 99.10 | 0.050 |
| 22.0 | 99.37 | 99.43 | 0.032 |
| 25.0 | 99.68 | 99.72 | 0.016 |
| 27.0 | 99.80 | 99.82 | 0.010 |
| 30.0 | 99.90 | 99.91 | 0.005 |
| 32.2 | 99.94 | 99.95 | 0.005 |
| 34.0 | 99.96 | 99.97 | 0.005 |

**Supplementary Table S2.** Precision in the longest overlap subset in the ul-hor-QV25 sample. Comparisons were made for chromosome 1 and chromosome 7 between mm2-ivh-hpc with $W = 3$ and mm2-k-hpc with $k = 133$. All values are rounded to three significant figures.

| Chromosome | Configuration | Precision in the longest overlap subset (%) |
|---|---|---|
| chr1 | mm2-ivh-hpc ($W = 3$) | 99.2 |
| | mm2-k-hpc ($k = 133$) | 29.7 |
| chr7 | mm2-ivh-hpc ($W = 3$) | 71.2 |
| | mm2-k-hpc ($k = 133$) | 17.8 |

**Supplementary Table S3.** Number of reads and total base count per chromosome (assembly bin) after binning the reads mapped to CHM13 v2.0 with minimap2.

| **Chromosome (Assembly bin)** | **Number of reads** | **Total base count (bp)** |
|---|---:|---:|
| chr1 | 1,879 | 236,969,559 |
| chr2 | 674 | 79,969,803 |
| chr3 | 2,490 | 300,366,940 |
| chr4 | 1,434 | 169,075,573 |
| chr5 | 1,989 | 236,832,575 |
| chr6 | 1,450 | 167,712,649 |
| chr7 | 1,021 | 121,569,818 |
| chr8 | 871 | 102,826,647 |
| chr9 | 809 | 97,236,606 |
| chr10 | 789 | 94,269,799 |
| chr11 | 948 | 116,160,650 |
| chr12 | 846 | 102,664,447 |
| chr13 | 1,676 | 194,132,350 |
| chr14 | 1,450 | 178,455,204 |
| chr15 | 833 | 93,339,637 |
| chr16 | 751 | 87,767,610 |
| chr17 | 1,064 | 122,142,216 |
| chr18 | 1,428 | 165,464,847 |
| chr19 | 1,394 | 165,554,986 |
| chr20 | 1,491 | 178,887,680 |
| chr21 | 1,429 | 171,879,188 |
| chr22 | 2,068 | 248,700,944 |
| chrX | 449 | 52,780,941 |
| chrY | 172 | 20,475,948 |

**Supplementary Table S4.** CPU time in seconds consumed for the entire assembly process (overlap detection and unitig generation) per chromosome for each assembler. Chromosome-wise values are rounded to three significant figures, and the total values are rounded to four significant figures.

| Chromosome (Assembly bin) | mm2-ivh-hpc | mm2-k133-hpc | hifiasm-ont-r1 |
|---|---:|---:|---:|
| chr1 | 1,460.0 | 187,000 | 745.0 |
| chr2 | 215.0 | 12,100 | 448.0 |
| chr3 | 1,980.0 | 103,000 | 1,300.0 |
| chr4 | 1,370.0 | 77,600 | 340.0 |
| chr5 | 2,300.0 | 54,300 | 692.0 |
| chr6 | 4,570.0 | 216,000 | 701.0 |
| chr7 | 3,910.0 | 16,000 | 484.0 |
| chr8 | 2,520.0 | 72,800 | 287.0 |
| chr9 | 228.0 | 38,000 | 267.0 |
| chr10 | 369.0 | 38,300 | 175.0 |
| chr11 | 4,680.0 | 23,800 | 716.0 |
| chr12 | 1,090.0 | 38,300 | 255.0 |
| chr13 | 644.0 | 36,800 | 543.0 |
| chr14 | 1,690.0 | 2,870 | 643.0 |
| chr15 | 838.0 | 11,100 | 303.0 |
| chr16 | 871.0 | 46,100 | 257.0 |
| chr17 | 6,020.0 | 89,000 | 516.0 |
| chr18 | 1,400.0 | 93,600 | 662.0 |
| chr19 | 1,080.0 | 9,320 | 946.0 |
| chr20 | 892.0 | 25,700 | 466.0 |
| chr21 | 425.0 | 2,650 | 452.0 |
| chr22 | 1,710.0 | 4,280 | 1,010.0 |
| chrX | 2,110.0 | 24,500 | 220.0 |
| chrY | 39.1 | 150 | 44.7 |
| Total | 42,430.0 | 1,223,000 | 12,470.0 |

**Supplementary Table S5.** CPU time in seconds consumed for the overlap detection step for mm2-ivh-hpc and mm2-k133-hpc. Chromosome-wise values are rounded to three significant figures, and the total values are rounded to four significant figures.

| Chromosome (Assembly bin) | mm2-ivh-hpc | mm2-k133-hpc |
|---|---:|---:|
| chr1 | 1,460.0 | 186,000 |
| chr2 | 215.0 | 12,100 |
| chr3 | 1,980.0 | 103,000 |
| chr4 | 1,370.0 | 77,400 |
| chr5 | 2,300.0 | 53,900 |
| chr6 | 4,560.0 | 216,000 |
| chr7 | 3,910.0 | 15,900 |
| chr8 | 2,520.0 | 72,600 |
| chr9 | 228.0 | 37,900 |
| chr10 | 369.0 | 38,200 |
| chr11 | 4,680.0 | 23,700 |
| chr12 | 1,090.0 | 38,200 |
| chr13 | 642.0 | 36,800 |
| chr14 | 1,690.0 | 2,840 |
| chr15 | 837.0 | 11,000 |
| chr16 | 870.0 | 46,000 |
| chr17 | 6,020.0 | 89,000 |
| chr18 | 1,400.0 | 93,200 |
| chr19 | 1,080.0 | 9,260 |
| chr20 | 891.0 | 25,700 |
| chr21 | 425.0 | 2,620 |
| chr22 | 1,710.0 | 4,240 |
| chrX | 2,110.0 | 24,500 |
| chrY | 39.0 | 150 |
| Total | 42,410.0 | 1,220,000 |

**Supplementary Table S6.** Maximum memory usage in GiB for each assembler per chromosome. All values are rounded to three significant figures.

| Chromosome (Assembly bin) | mm2-ivh-hpc | mm2-k133-hpc | hifiasm-ont-r1 |
|---|---:|---:|---:|
| chr1 | 7.14 | 181.00 | 16.6 |
| chr2 | 1.82 | 27.20 | 16.3 |
| chr3 | 8.67 | 117.00 | 16.6 |
| chr4 | 7.69 | 68.20 | 16.6 |
| chr5 | 9.22 | 95.40 | 16.8 |
| chr6 | 12.90 | 133.00 | 16.6 |
| chr7 | 13.00 | 28.10 | 16.5 |
| chr8 | 9.43 | 68.40 | 16.4 |
| chr9 | 1.67 | 48.20 | 16.4 |
| chr10 | 3.24 | 45.50 | 16.5 |
| chr11 | 14.00 | 40.50 | 16.5 |
| chr12 | 7.15 | 37.20 | 16.5 |
| chr13 | 5.44 | 38.70 | 16.7 |
| chr14 | 7.98 | 14.60 | 16.6 |
| chr15 | 6.84 | 22.20 | 16.5 |
| chr16 | 6.31 | 56.40 | 16.3 |
| chr17 | 12.70 | 77.70 | 16.5 |
| chr18 | 7.71 | 88.40 | 16.6 |
| chr19 | 5.57 | 29.90 | 16.6 |
| chr20 | 6.24 | 32.30 | 16.6 |
| chr21 | 3.46 | 11.30 | 16.6 |
| chr22 | 9.49 | 16.20 | 16.7 |
| chrX | 9.61 | 29.50 | 16.2 |
| chrY | 1.84 | 4.61 | 16.2 |
| Geometric mean | 6.43 | 40.50 | 16.5 |

**Supplementary Table S7.** Sizes in GiB of the output PAF files from the overlap detection step for mm2-ivh-hpc and mm2-k133-hpc, shown per chromosome. Chromosome-wise values are rounded to three significant figures, and the total values are rounded to four significant figures.

| Chromosome (Assembly bin) | mm2-ivh-hpc | mm2-k133-hpc |
|---|---:|---:|
| chr1 | 2.400 | 194.000 |
| chr2 | 0.547 | 24.800 |
| chr3 | 4.800 | 134.000 |
| chr4 | 2.190 | 68.800 |
| chr5 | 5.510 | 139.000 |
| chr6 | 7.250 | 57.800 |
| chr7 | 8.560 | 29.200 |
| chr8 | 3.010 | 34.800 |
| chr9 | 0.536 | 39.600 |
| chr10 | 0.804 | 43.700 |
| chr11 | 11.200 | 40.600 |
| chr12 | 3.390 | 25.000 |
| chr13 | 1.330 | 30.700 |
| chr14 | 4.050 | 13.400 |
| chr15 | 0.670 | 9.550 |
| chr16 | 2.390 | 17.900 |
| chr17 | 3.810 | 20.600 |
| chr18 | 3.220 | 88.300 |
| chr19 | 2.570 | 38.600 |
| chr20 | 2.140 | 20.000 |
| chr21 | 1.150 | 8.890 |
| chr22 | 4.210 | 15.800 |
| chrX | 1.220 | 5.730 |
| chrY | 0.013 | 0.169 |
| Total | 76.990 | 1,101.000 |

**Supplementary Table S8.** Visual evaluation results of the assembly. Haplotypes where a single unitig covered the entire length of the corresponding HG002 v1.1 active HOR region were marked as complete. Haplotypes where a single unitig covered approximately 90% or more of the active HOR region were classified as near-complete. Haplotypes that covered the entire length of the active HOR region by tiling multiple unitigs (regardless of the presence of correct edges between them) were marked as split. Haplotypes that included unitigs with inconsistent mappings to the entire or part of the active HOR region were marked as collapse. Haplotypes for which no unitig corresponding to any interval of the active HOR region was generated were marked as missing.

| **Chromosome (Assembly bin)** | **Haplotype** | **mm2-ivh-hpc** | **mm2-k133-hpc** | **hifiasm-ont-r1** |
|---|---|---:|---:|---:|
| chr1 | maternal | **complete** | collapse | **complete** |
| | paternal | **complete** | collapse | **complete** |
| chr2 | maternal | **complete** | collapse | **complete** |
| | paternal | **complete** | collapse | **complete** |
| chr3 | maternal | split | collapse | split |
| | paternal | collapse | collapse | collapse |
| chr4 | maternal | **complete** | collapse | missing |
| | paternal | near-complete | collapse | **complete** |
| chr5 | maternal | **complete** | missing | **complete** |
| | paternal | collapse | missing | collapse |
| chr6 | maternal | near-complete | collapse | collapse |
| | paternal | near-complete | missing | collapse |
| chr7 | maternal | **complete** | missing | collapse |
| | paternal | split | missing | collapse |
| chr8 | maternal | near-complete | collapse | **complete** |
| | paternal | **complete** | missing | missing |
| chr9 | maternal | **complete** | collapse | missing |
| | paternal | **complete** | missing | **complete** |
| chr10 | maternal | split | collapse | split |
| | paternal | **complete** | missing | **complete** |
| chr11 | maternal | **complete** | collapse | collapse |
| | paternal | split | collapse | collapse |
| chr12 | maternal | **complete** | missing | collapse |
| | paternal | **complete** | missing | missing |
| chr13 | maternal | **complete** | missing | **complete** |
| | paternal | missing | missing | missing |
| chr14 | maternal | split | collapse | missing |
| | paternal | **complete** | missing | collapse |
| chr15 | maternal | **complete** | collapse | missing |
| | paternal | **complete** | missing | **complete** |
| chr16 | maternal | split | missing | missing |
| | paternal | **complete** | missing | **complete** |
| chr17 | maternal | **complete** | collapse | collapse |
| | paternal | split | collapse | collapse |
| chr18 | maternal | split | collapse | collapse |
| | paternal | split | missing | **complete** |
| chr19 | maternal | **complete** | collapse | collapse |
| | paternal | split | collapse | collapse |
| chr20 | maternal | split | collapse | **complete** |
| | paternal | split | collapse | **complete** |
| chr21 | maternal | **complete** | collapse | collapse |
| | paternal | **complete** | missing | collapse |
| chr22 | maternal | **complete** | collapse | collapse |
| | paternal | **complete** | collapse | collapse |
| chrX | | **complete** | collapse | collapse |
| chrY | | **complete** | **complete** | **complete** |

**Supplementary Table S9.** Reads sampled around the misjoin point in the Hifiasm output

| Name in Figure 1(e) | Original read ID | Read length | Mapped coordinates |
|---|---|---|---|
| mat1 | ef3b6bf7-a289-49d1-944a-271a8741296f | 294,291 | chr21_MATERNAL:13,074,863-13,368,848 |
| mat2 | 89f68e08-13fd-423b-b5b8-29407939fd25 | 244,376 | chr21_MATERNAL:13,008,260-13,252,592 |
| pat1 | 637b9c25-af68-461c-bdcf-5d277ba4061e | 261,159 | chr21_PATERNAL:9,528,117-9,788,970 |
| pat2 | efac66f0-07f2-4dca-9c77-3acd6d47aa1d | 278,347 | chr21_PATERNAL:9,476,531-9,754,673 |

**Supplementary Table S10.** The sample-matched reference version of Supplementary Table S3. Number of reads and total base count per chromosome (assembly bin) after binning the reads mapped to HG002 v1.1 with minimap2.

| Chromosome (Assembly bin) | Number of reads | Total base count (bp) |
|---|---:|---:|
| chr1 | 1,463 | 177,774,794 |
| chr2 | 674 | 79,969,803 |
| chr3 | 2,063 | 240,361,703 |
| chr4 | 1,707 | 201,924,780 |
| chr5 | 1,956 | 230,863,550 |
| chr6 | 1,446 | 167,118,221 |
| chr7 | 987 | 117,671,378 |
| chr8 | 872 | 102,965,235 |
| chr9 | 809 | 97,236,606 |
| chr10 | 794 | 94,690,510 |
| chr11 | 944 | 114,893,916 |
| chr12 | 842 | 101,748,162 |
| chr13 | 2,006 | 234,398,094 |
| chr14 | 1,857 | 223,530,858 |
| chr15 | 1,209 | 138,882,730 |
| chr16 | 775 | 89,784,096 |
| chr17 | 1,072 | 122,860,117 |
| chr18 | 1,429 | 165,514,953 |
| chr19 | 1,338 | 156,720,172 |
| chr20 | 1,481 | 177,000,577 |
| chr21 | 1,144 | 132,844,338 |
| chr22 | 1,736 | 207,954,706 |
| chrX | 449 | 52,780,941 |
| chrY | 171 | 20,259,101 |

**Supplementary Table S11.** The sample-matched reference version of Supplementary Table S4. CPU time in seconds consumed for the entire assembly process (overlap detection and unitig generation) per chromosome for each assembler. Chromosome-wise values are rounded to three significant figures, and the total values are rounded to four significant figures.

| Chromosome (Assembly bin) | mm2-ivh-hpc | mm2-k133-hpc | hifiasm-ont-r1 |
|---|---:|---:|---:|
| chr1 | 815.0 | 79,800 | 734.0 |
| chr2 | 217.0 | 10,500 | 436.0 |
| chr3 | 1,870.0 | 92,600 | 968.0 |
| chr4 | 1,300.0 | 50,700 | 691.0 |
| chr5 | 2,350.0 | 61,600 | 651.0 |
| chr6 | 4,960.0 | 204,000 | 655.0 |
| chr7 | 4,090.0 | 18,600 | 325.0 |
| chr8 | 2,670.0 | 60,300 | 323.0 |
| chr9 | 245.0 | 32,400 | 254.0 |
| chr10 | 400.0 | 38,500 | 218.0 |
| chr11 | 4,780.0 | 27,000 | 669.0 |
| chr12 | 1,130.0 | 37,300 | 246.0 |
| chr13 | 789.0 | 38,600 | 576.0 |
| chr14 | 1,720.0 | 1,230 | 651.0 |
| chr15 | 1,070.0 | 9,440 | 428.0 |
| chr16 | 924.0 | 40,300 | 253.0 |
| chr17 | 6,380.0 | 82,200 | 516.0 |
| chr18 | 1,450.0 | 86,100 | 643.0 |
| chr19 | 1,170.0 | 8,910 | 910.0 |
| chr20 | 933.0 | 28,000 | 438.0 |
| chr21 | 285.0 | 1,100 | 289.0 |
| chr22 | 1,700.0 | 4,610 | 602.0 |
| chrX | 2,230.0 | 22,000 | 207.0 |
| chrY | 37.7 | 125 | 43.2 |
| Total | 43,530.0 | 1,036,000 | 11,720.0 |

**Supplementary Table S12.** The sample-matched reference version of Supplementary Table S5. CPU time in seconds consumed for the overlap detection step for mm2-ivh and mm2-k133. Chromosome-wise values are rounded to three significant figures, and the total values are rounded to four significant figures.

| Chromosome (Assembly bin) | mm2-ivh-hpc | mm2-k133-hpc |
|---|---:|---:|
| chr1 | 815.0 | 79,400 |
| chr2 | 217.0 | 10,400 |
| chr3 | 1,870.0 | 92,400 |
| chr4 | 1,300.0 | 50,600 |
| chr5 | 2,350.0 | 61,300 |
| chr6 | 4,950.0 | 204,000 |
| chr7 | 4,090.0 | 18,500 |
| chr8 | 2,670.0 | 60,200 |
| chr9 | 245.0 | 32,300 |
| chr10 | 399.0 | 38,300 |
| chr11 | 4,780.0 | 26,800 |
| chr12 | 1,130.0 | 37,300 |
| chr13 | 788.0 | 38,500 |
| chr14 | 1,720.0 | 1,210 |
| chr15 | 1,070.0 | 9,420 |
| chr16 | 923.0 | 40,300 |
| chr17 | 6,380.0 | 82,100 |
| chr18 | 1,450.0 | 85,900 |
| chr19 | 1,170.0 | 8,850 |
| chr20 | 932.0 | 28,000 |
| chr21 | 284.0 | 1,090 |
| chr22 | 1,700.0 | 4,590 |
| chrX | 2,230.0 | 22,000 |
| chrY | 37.7 | 124 |
| Total | 43,510.0 | 1,033,000 |

**Supplementary Table S13.** The sample-matched reference version of Supplementary Table S6. Maximum memory usage in GiB for each assembler per chromosome. All values are rounded to three significant figures.

| Chromosome (Assembly bin) | mm2-ivh-hpc | mm2-k133-hpc | hifiasm-ont-r1 |
|---|---|---|---|
| chr1 | 2.68 | 106.00 | 16.6 |
| chr2 | 1.83 | 27.60 | 16.3 |
| chr3 | 8.55 | 94.50 | 16.6 |
| chr4 | 7.29 | 51.10 | 16.5 |
| chr5 | 9.12 | 110.00 | 16.6 |
| chr6 | 13.80 | 137.00 | 16.6 |
| chr7 | 12.70 | 32.00 | 16.4 |
| chr8 | 10.30 | 70.50 | 16.4 |
| chr9 | 1.73 | 48.30 | 16.4 |
| chr10 | 3.13 | 47.70 | 16.3 |
| chr11 | 14.60 | 45.00 | 16.3 |
| chr12 | 7.21 | 38.50 | 16.4 |
| chr13 | 5.46 | 42.60 | 16.7 |
| chr14 | 8.71 | 13.40 | 16.6 |
| chr15 | 7.10 | 17.80 | 16.5 |
| chr16 | 6.35 | 49.80 | 16.4 |
| chr17 | 13.40 | 75.60 | 16.4 |
| chr18 | 8.01 | 86.70 | 16.6 |
| chr19 | 5.93 | 31.60 | 16.6 |
| chr20 | 6.25 | 34.80 | 16.6 |
| chr21 | 2.91 | 8.47 | 16.5 |
| chr22 | 8.90 | 18.70 | 16.6 |
| chrX | 9.05 | 30.10 | 16.2 |
| chrY | 1.85 | 4.63 | 16.2 |
| Geometric mean | 6.21 | 39.00 | 16.5 |

**Supplementary Table S14.** The sample-matched reference version of Supplementary Table S7. Sizes in GiB of the output PAF files from the overlap detection step for mm2-ivh-hpc and mm2-k133-hpc, shown per chromosome. Chromosome-wise values are rounded to three significant figures, and the total values are rounded to four significant figures.

| Chromosome (Assembly bin) | mm2-ivh-hpc | mm2-k133-hpc |
|---|---:|---:|
| chr1 | 1.070 | 130.000 |
| chr2 | 0.547 | 24.800 |
| chr3 | 4.320 | 104.000 |
| chr4 | 2.130 | 53.500 |
| chr5 | 5.560 | 156.000 |
| chr6 | 7.280 | 65.100 |
| chr7 | 8.550 | 30.900 |
| chr8 | 3.010 | 34.800 |
| chr9 | 0.536 | 39.600 |
| chr10 | 0.806 | 44.400 |
| chr11 | 11.300 | 43.700 |
| chr12 | 3.410 | 25.100 |
| chr13 | 1.690 | 39.800 |
| chr14 | 4.150 | 13.500 |
| chr15 | 1.050 | 11.100 |
| chr16 | 2.410 | 16.400 |
| chr17 | 3.810 | 20.600 |
| chr18 | 3.220 | 88.300 |
| chr19 | 2.840 | 42.400 |
| chr20 | 2.140 | 20.300 |
| chr21 | 0.705 | 4.470 |
| chr22 | 3.950 | 15.300 |
| chrX | 1.220 | 5.730 |
| chrY | 0.012 | 0.166 |
| Total | 75.680 | 1,030.000 |

**Supplementary Table S15.** The sample-matched reference version of Supplementary Table S8. The number of overlap records in the output PAF files from the overlap detection step for mm2-ivh-hpc and mm2-k133-hpc, shown per chromosome.

| Chromosome (Assembly bin) | Haplotype | mm2-ivh-hpc | mm2-k133-hpc | hifiasm-ont-r1 |
|---|---|---|---|---|
| chr1 | maternal | **complete** | collapse | missing |
| | paternal | **complete** | collapse | **complete** |
| chr2 | maternal | **complete** | collapse | **complete** |
| | paternal | **complete** | collapse | **complete** |
| chr3 | maternal | split | collapse | collapse |
| | paternal | collapse | collapse | collapse |
| chr4 | maternal | split | collapse | missing |
| | paternal | near-complete | collapse | **complete** |
| chr5 | maternal | **complete** | missing | **complete** |
| | paternal | collapse | missing | collapse |
| chr6 | maternal | near-complete | collapse | collapse |
| | paternal | near-complete | missing | collapse |
| chr7 | maternal | near-complete | collapse | collapse |
| | paternal | split | missing | collapse |
| chr8 | maternal | near-complete | collapse | collapse |
| | paternal | **complete** | missing | missing |
| chr9 | maternal | **complete** | collapse | missing |
| | paternal | **complete** | missing | **complete** |
| chr10 | maternal | split | collapse | split |
| | paternal | **complete** | collapse | **complete** |
| chr11 | maternal | near-complete | collapse | collapse |
| | paternal | split | collapse | collapse |
| chr12 | maternal | **complete** | missing | collapse |
| | paternal | **complete** | missing | near-complete |
| chr13 | maternal | **complete** | collapse | missing |
| | paternal | **complete** | collapse | **complete** |
| chr14 | maternal | collapse | collapse | **complete** |
| | paternal | **complete** | split | missing |
| chr15 | maternal | **complete** | collapse | **complete** |
| | paternal | **complete** | missing | missing |
| chr16 | maternal | split | missing | missing |
| | paternal | **complete** | missing | **complete** |
| chr17 | maternal | near-complete | collapse | collapse |
| | paternal | split | collapse | collapse |
| chr18 | maternal | split | collapse | collapse |
| | paternal | split | missing | collapse |
| chr19 | maternal | **complete** | collapse | collapse |
| | paternal | split | collapse | collapse |
| chr20 | maternal | split | collapse | collapse |
| | paternal | split | collapse | collapse |
| chr21 | maternal | **complete** | collapse | **complete** |
| | paternal | **complete** | missing | **complete** |
| chr22 | maternal | **complete** | collapse | missing |
| | paternal | **complete** | collapse | collapse |
| chrX | | **complete** | collapse | collapse |
| chrY | | **complete** | **complete** | **complete** |
