## Supplementary figures and images for "mm2-ivh: simple and precise overlap detection in alpha satellite HORs with interval hashing"

### hifiasm-ont-r1.png

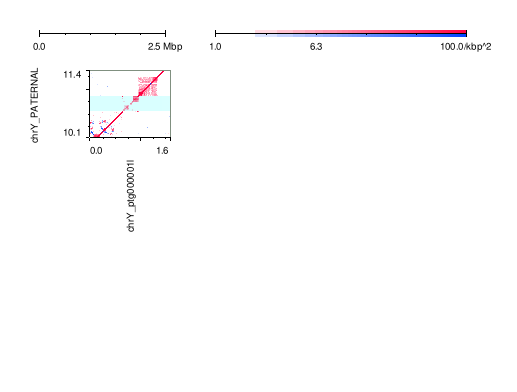

### hifiasm-ont-r1.png

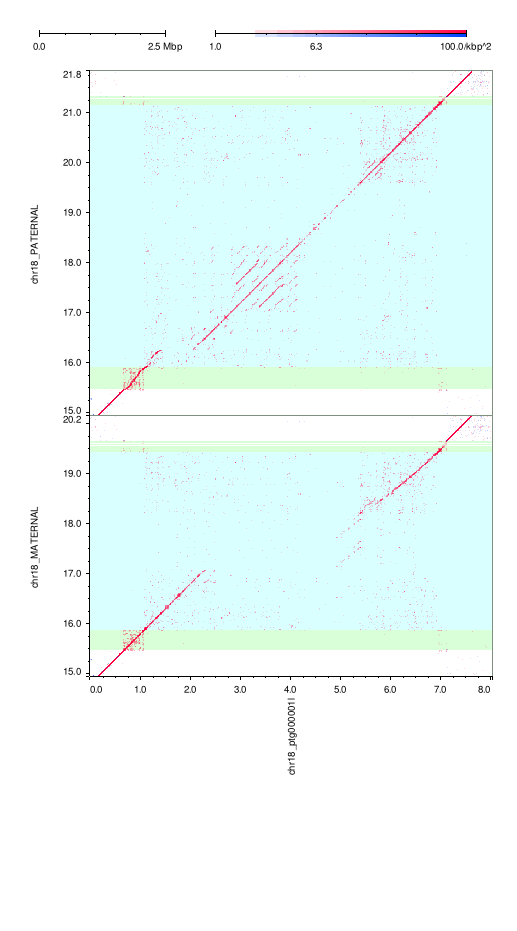

### hifiasm-ont-r1.png

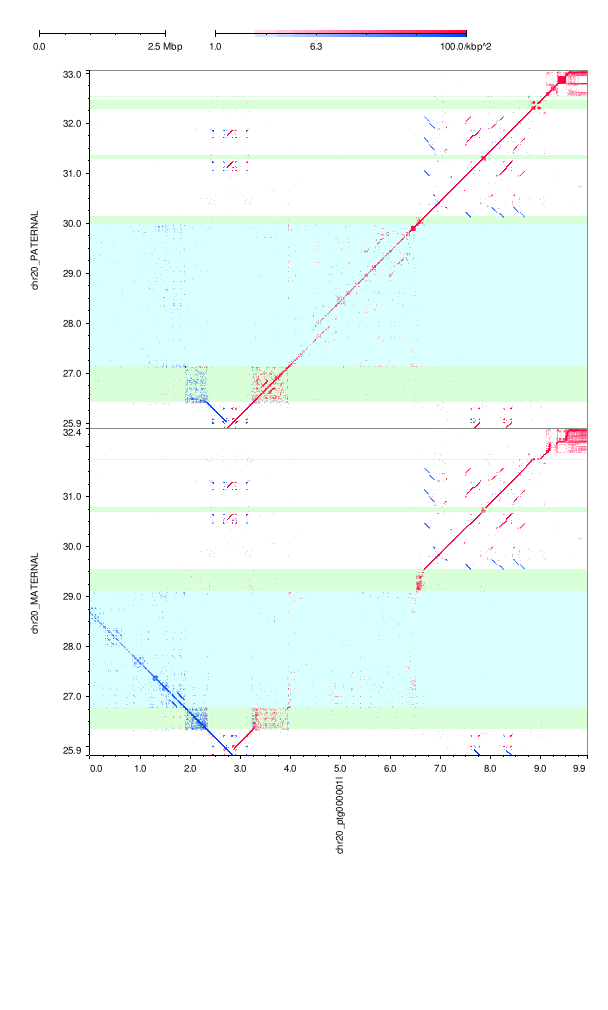

### hifiasm-ont-r1.png

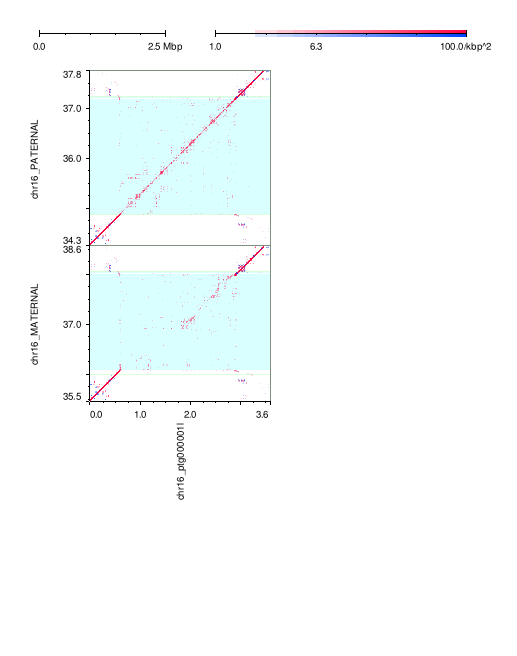

### hifiasm-ont-r1.png

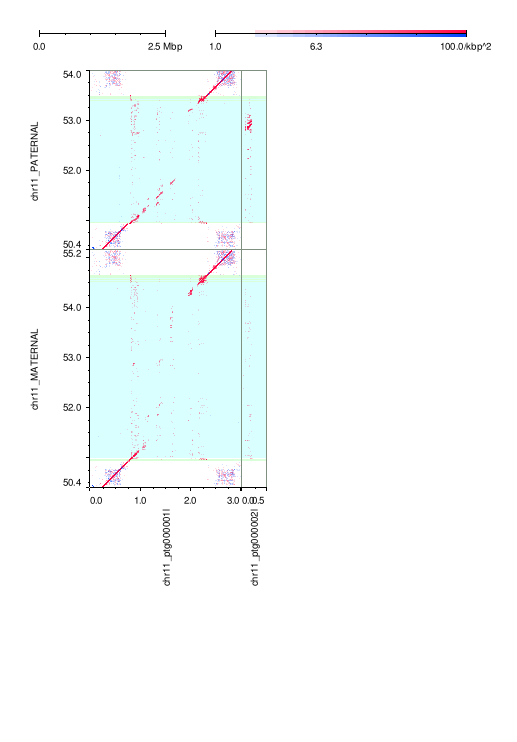

### hifiasm-ont-r1.png

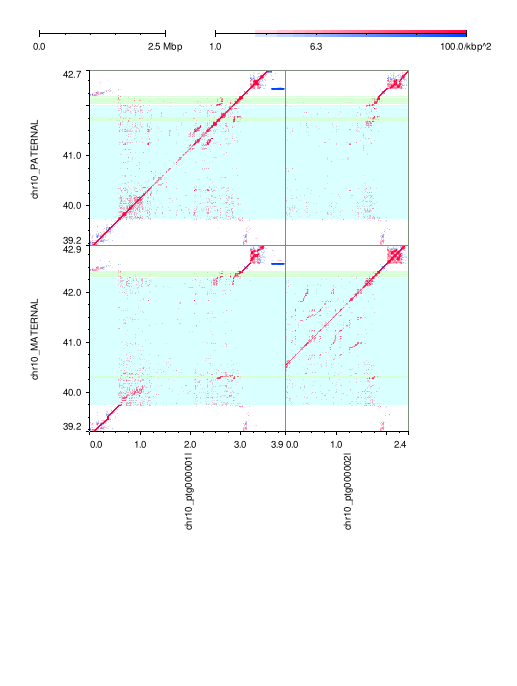

### hifiasm-ont-r1.png

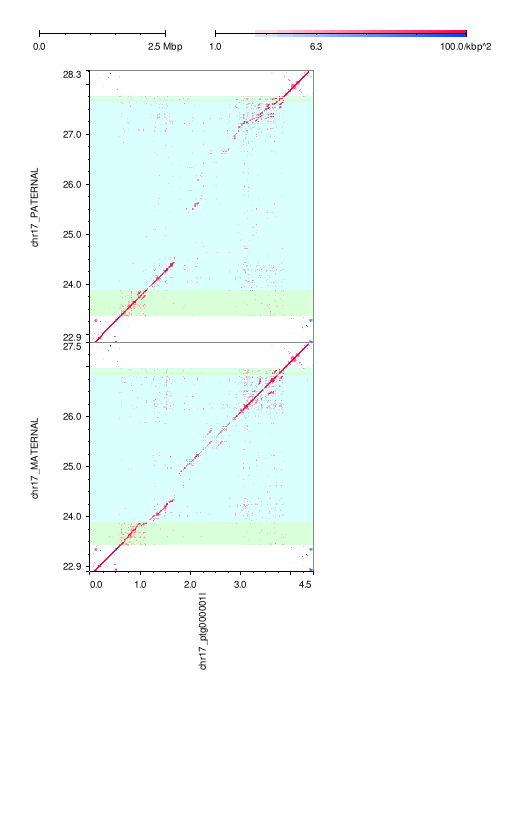

### hifiasm-ont-r1.png

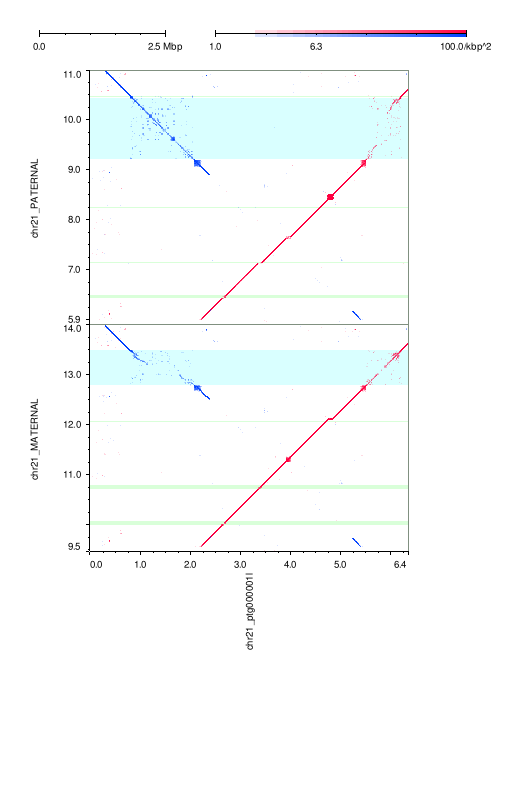

### hifiasm-ont-r1.png

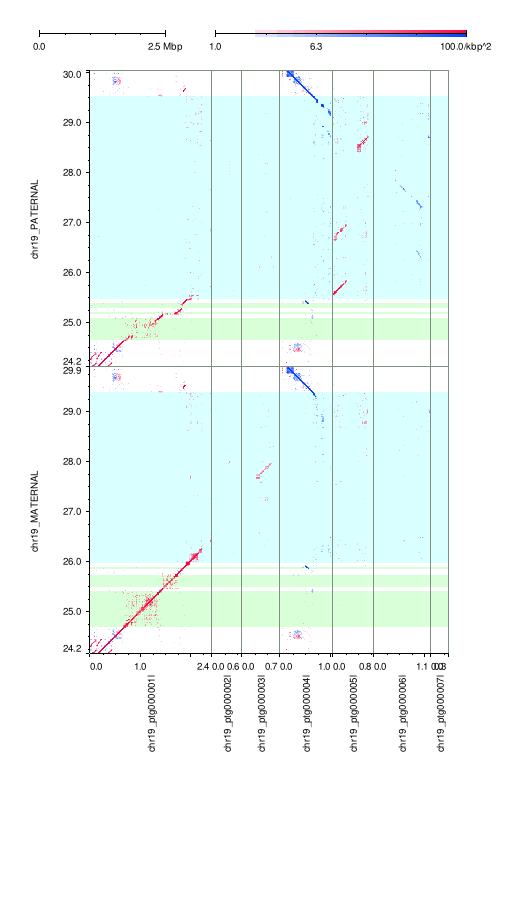

### hifiasm-ont-r1.png

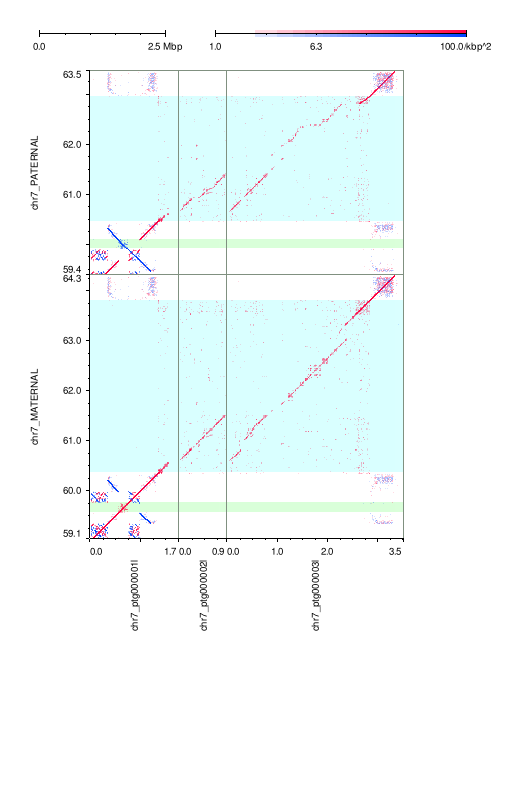

### hifiasm-ont-r1.png

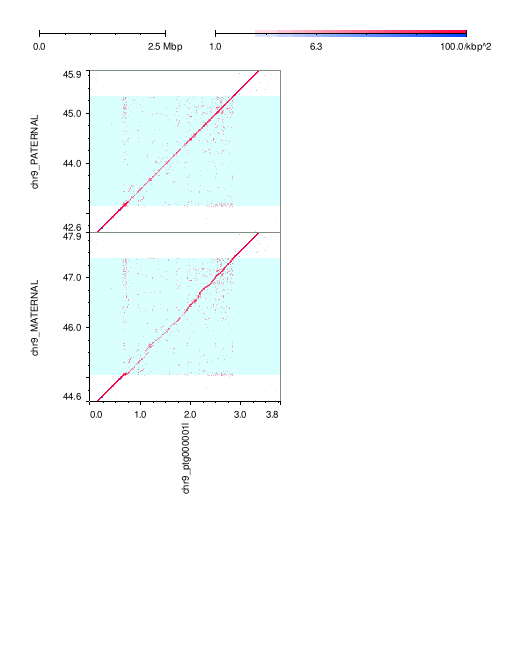

### hifiasm-ont-r1.png

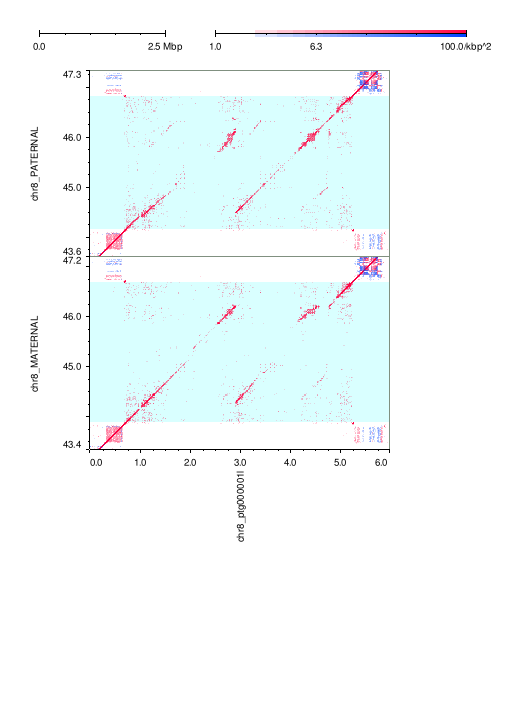

### hifiasm-ont-r1.png

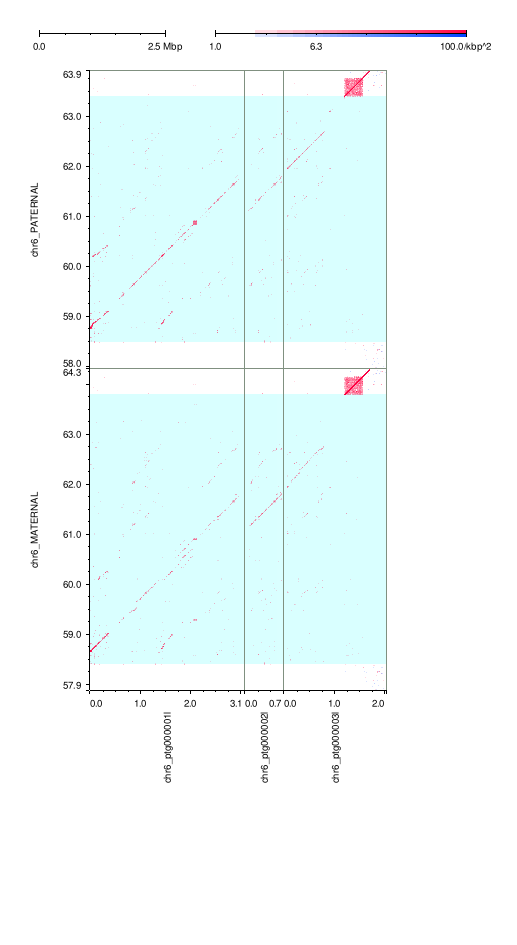

### hifiasm-ont-r1.png

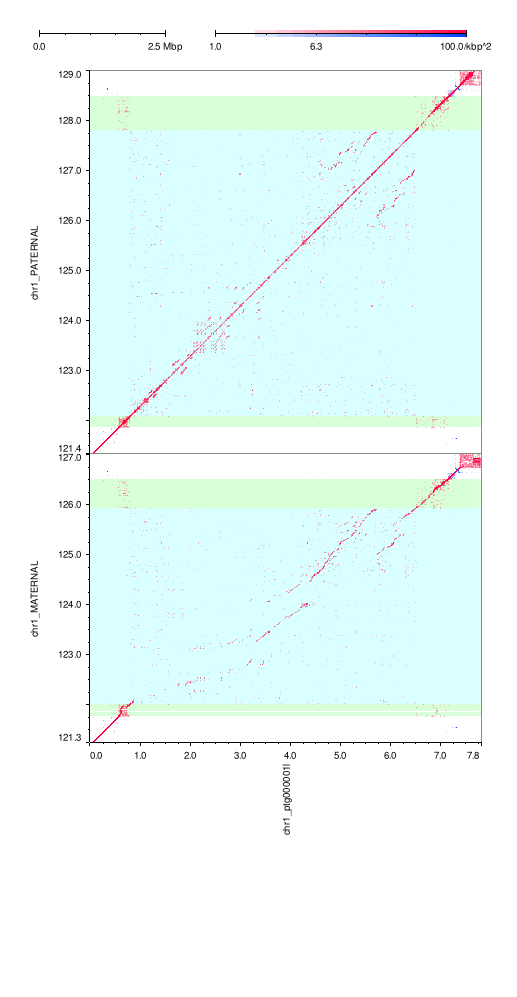

### mm2-ivh-hpc.png

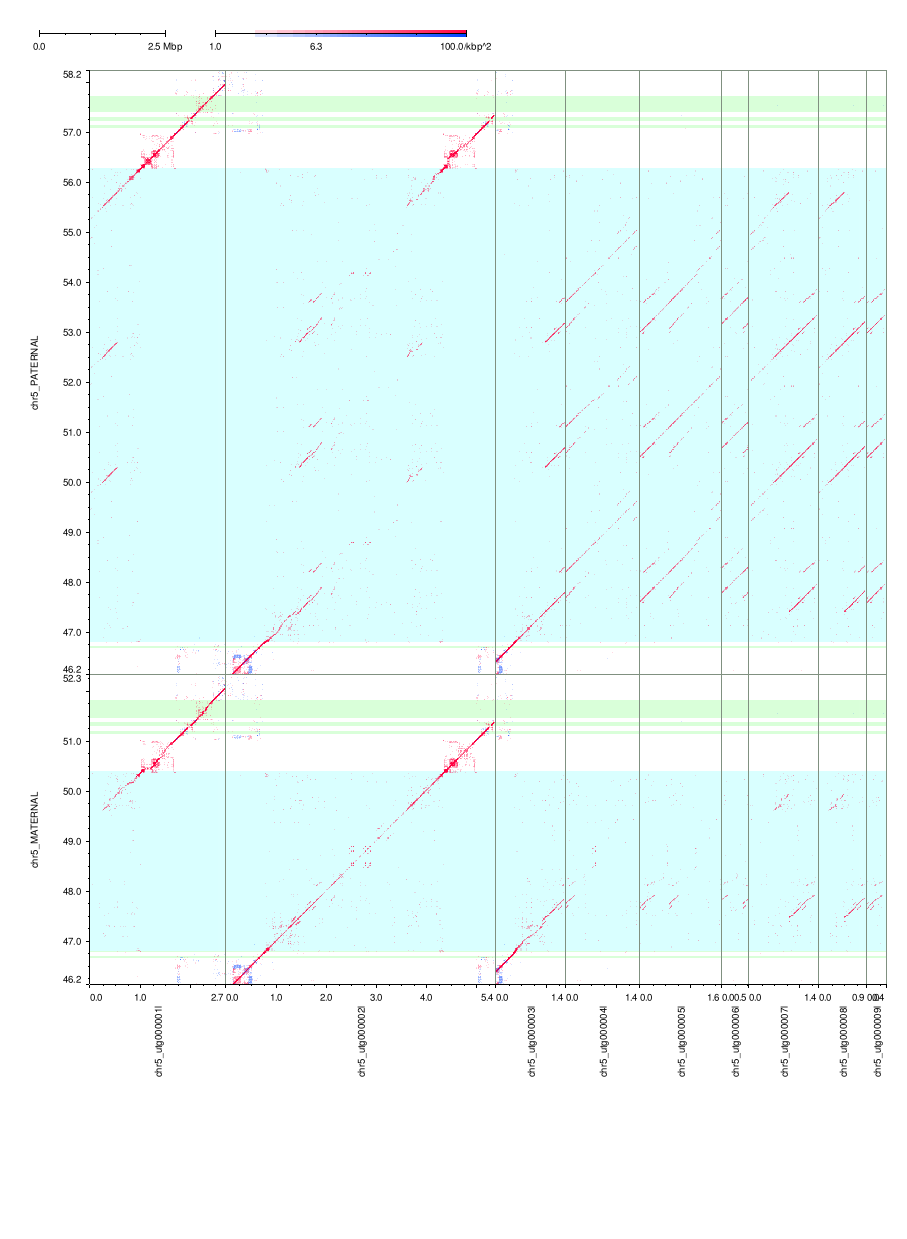

### mm2-ivh-hpc.png

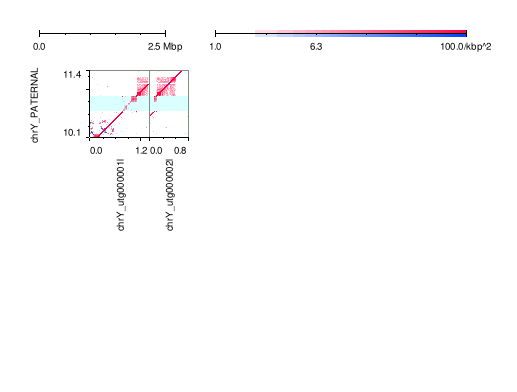

### mm2-ivh-hpc.png

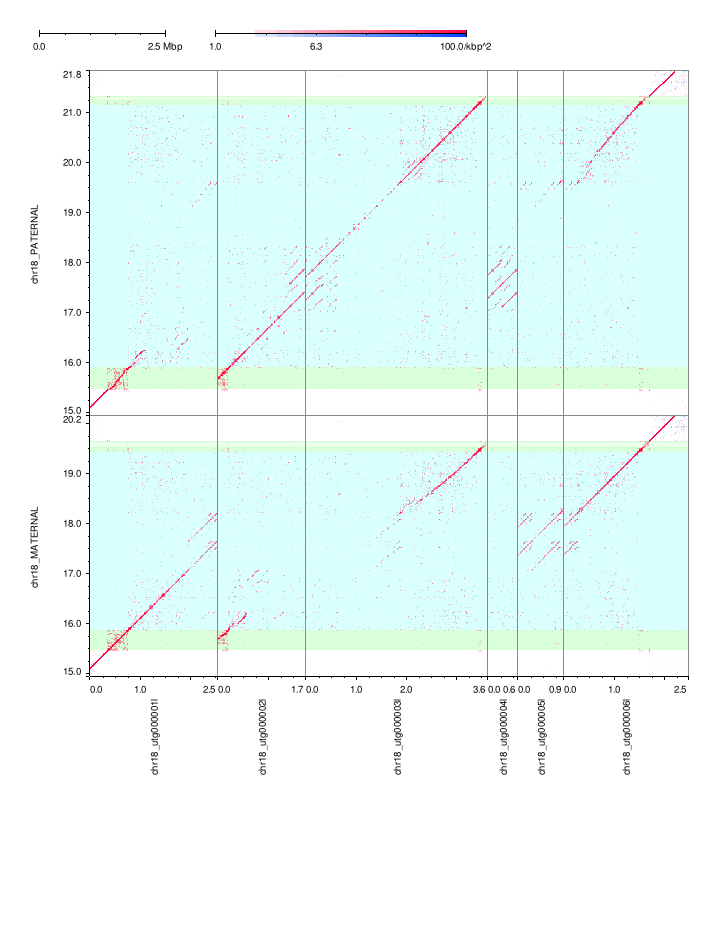

### mm2-ivh-hpc.png

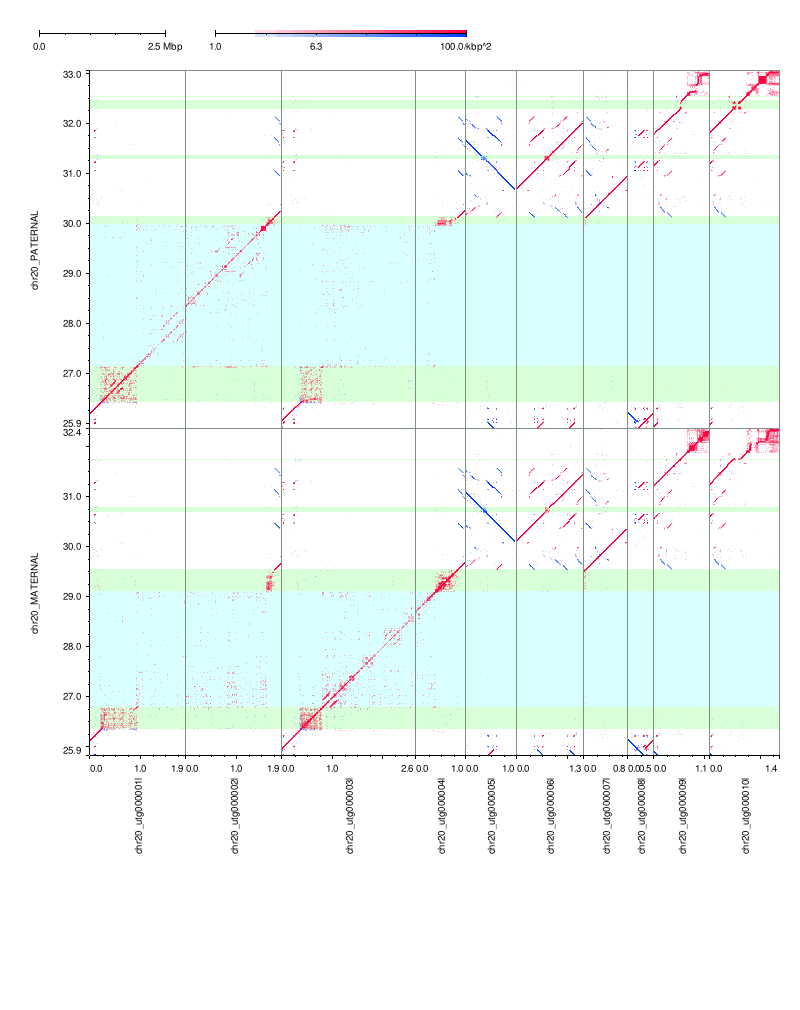

### mm2-ivh-hpc.png

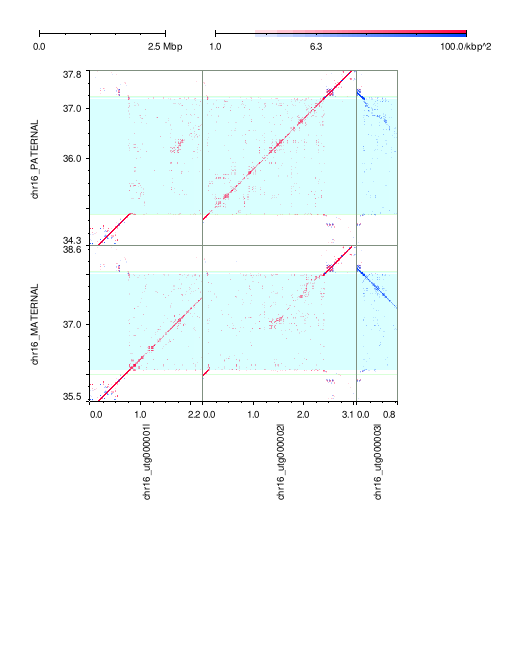

### mm2-ivh-hpc.png

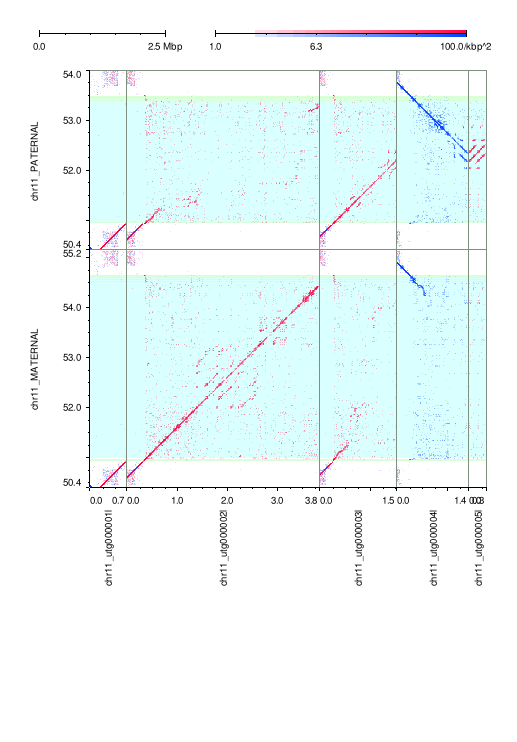

### mm2-ivh-hpc.png

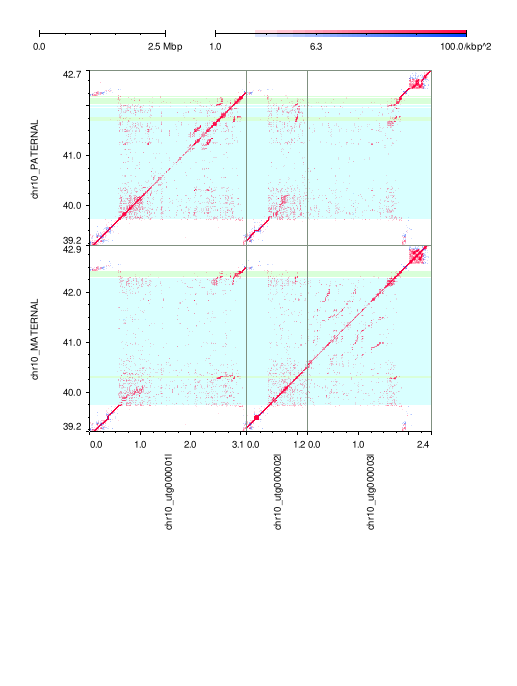

### mm2-ivh-hpc.png

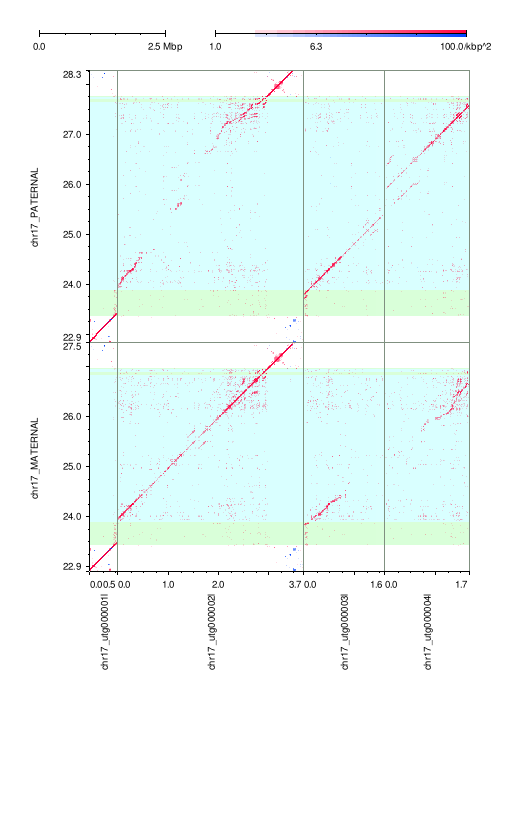

### mm2-ivh-hpc.png

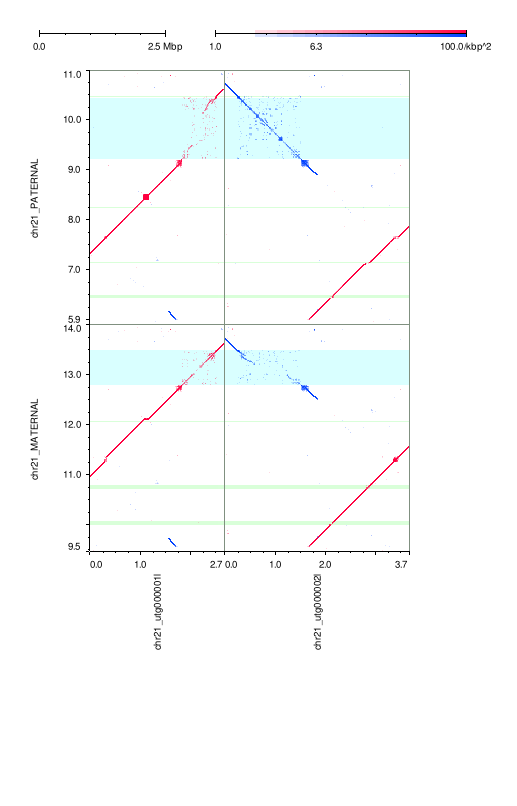

### mm2-ivh-hpc.png

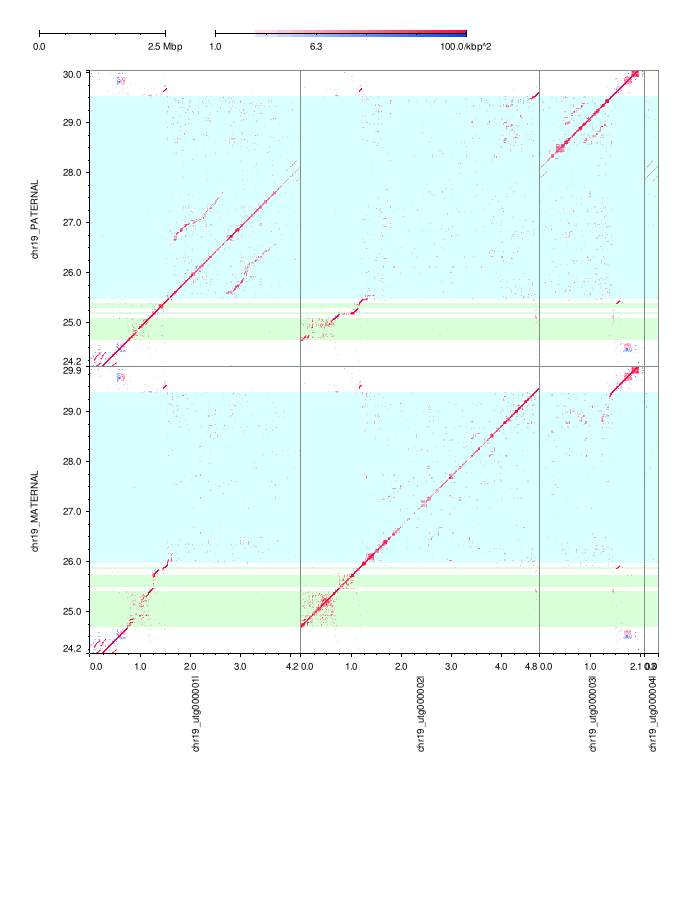

### mm2-ivh-hpc.png

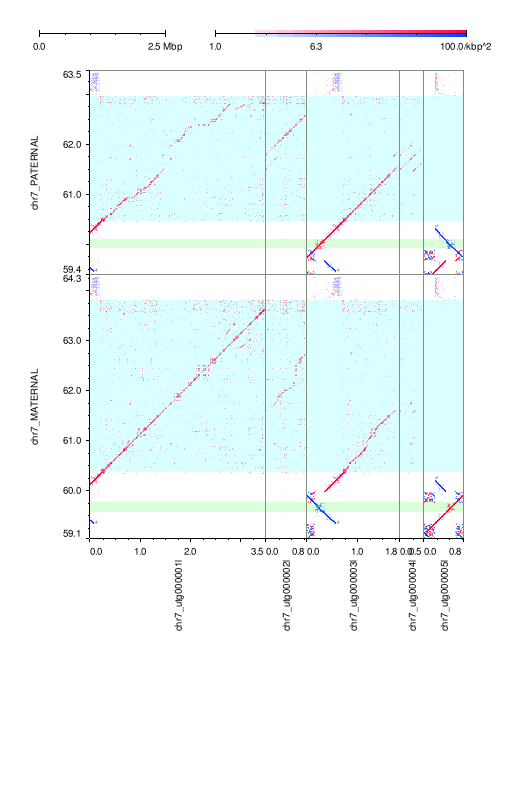

### mm2-ivh-hpc.png

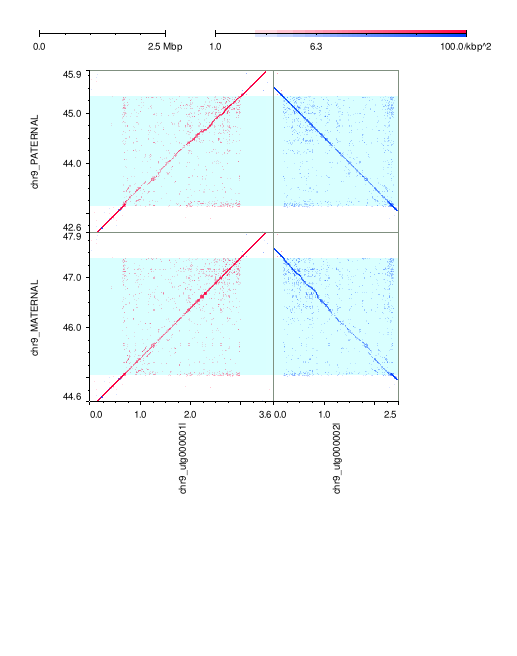

### mm2-ivh-hpc.png

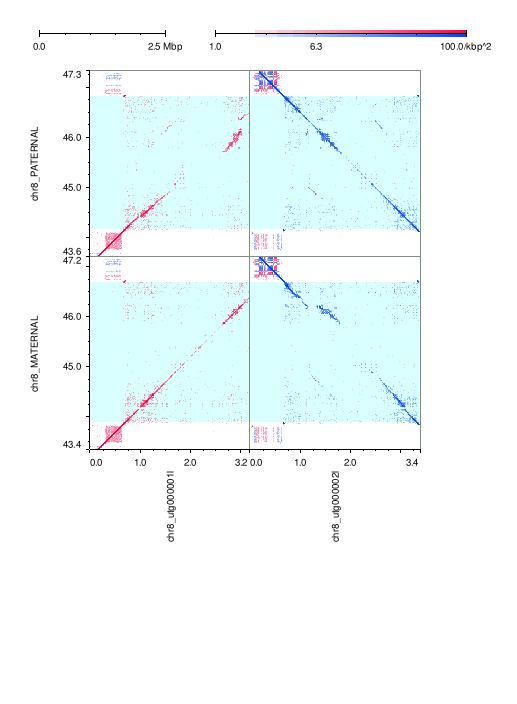

### mm2-ivh-hpc.png

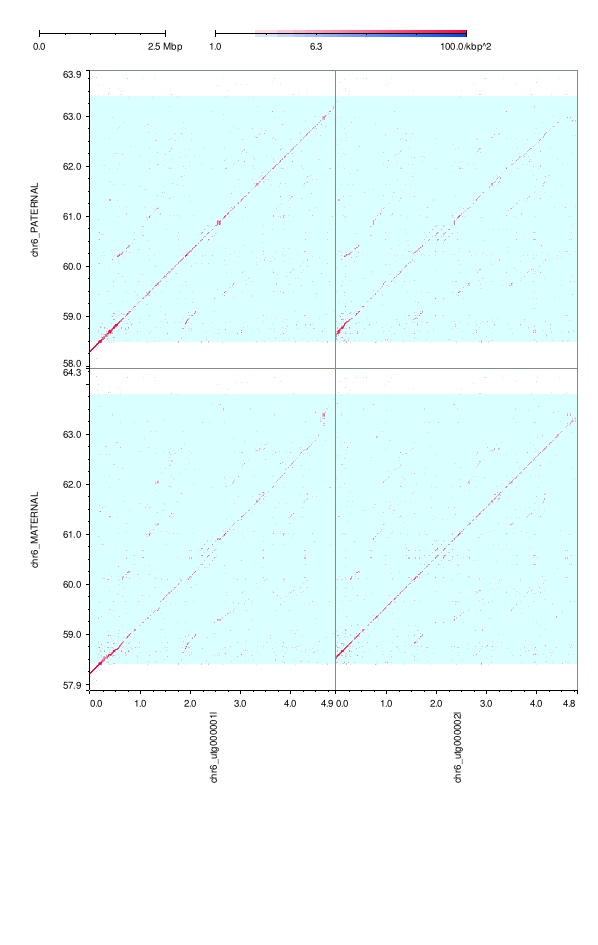

### mm2-ivh-hpc.png

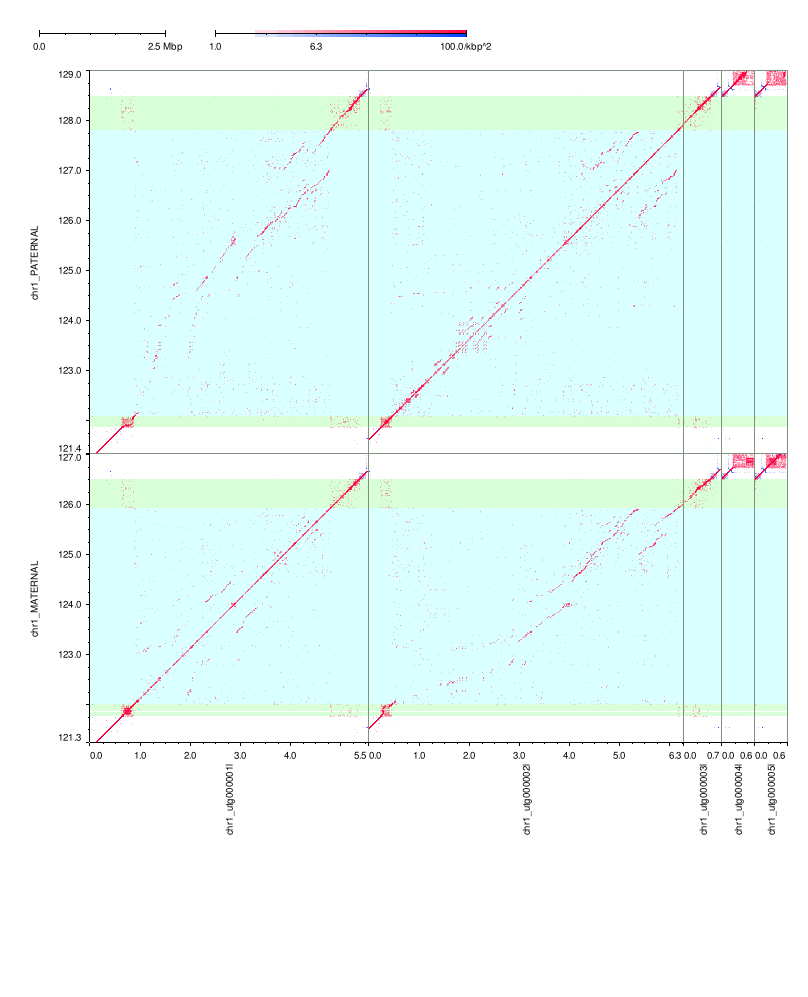

### mm2-k133-hpc.png

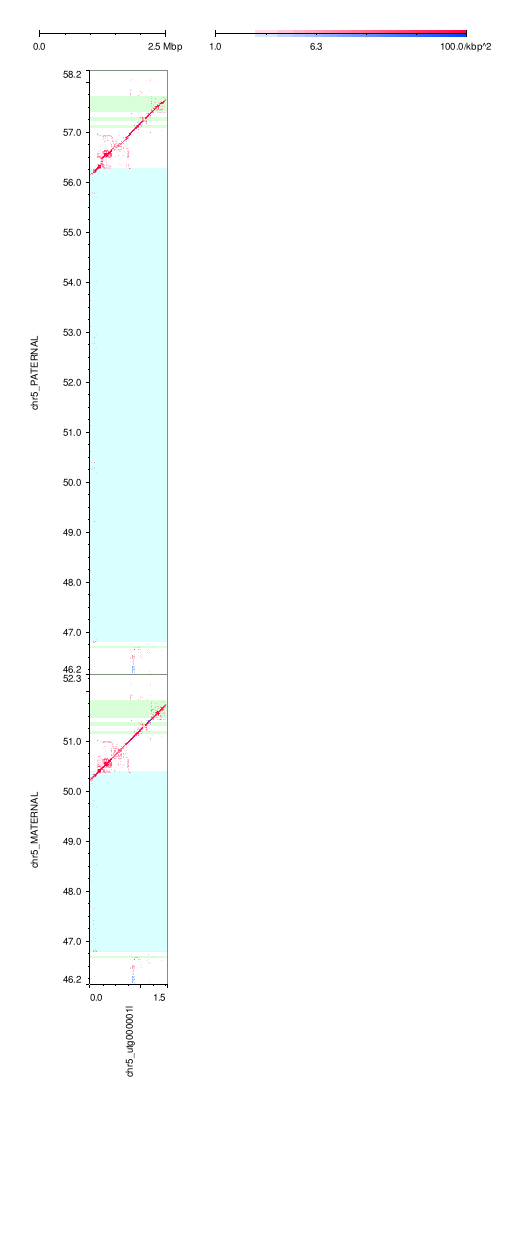

### mm2-k133-hpc.png

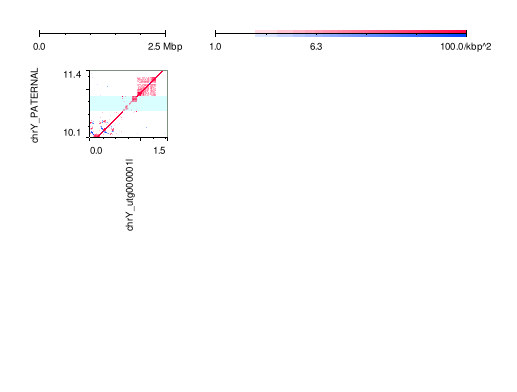

### mm2-k133-hpc.png

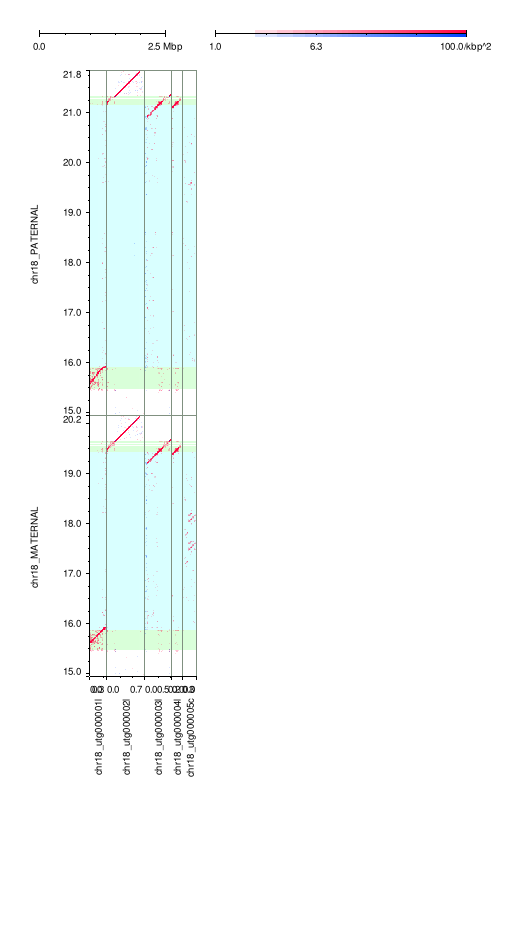

### mm2-k133-hpc.png

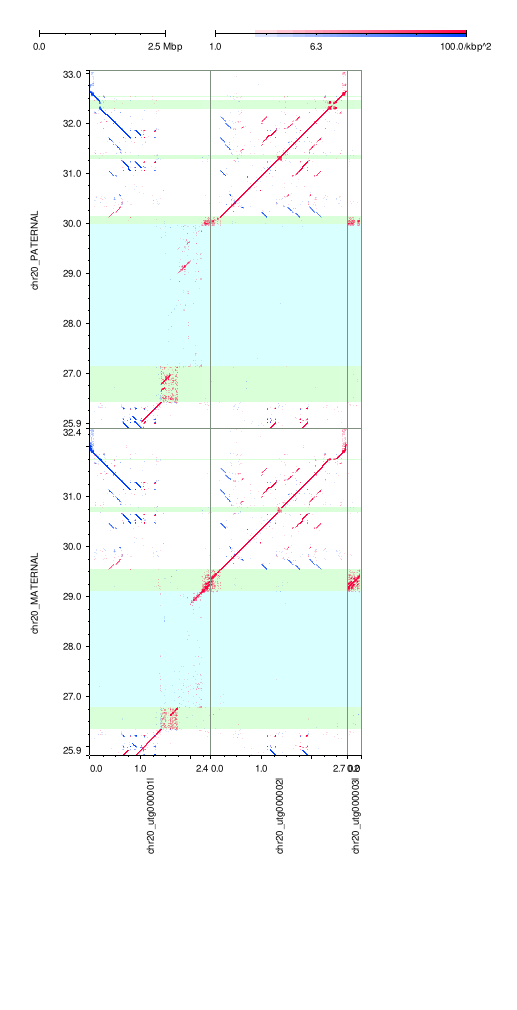

### mm2-k133-hpc.png

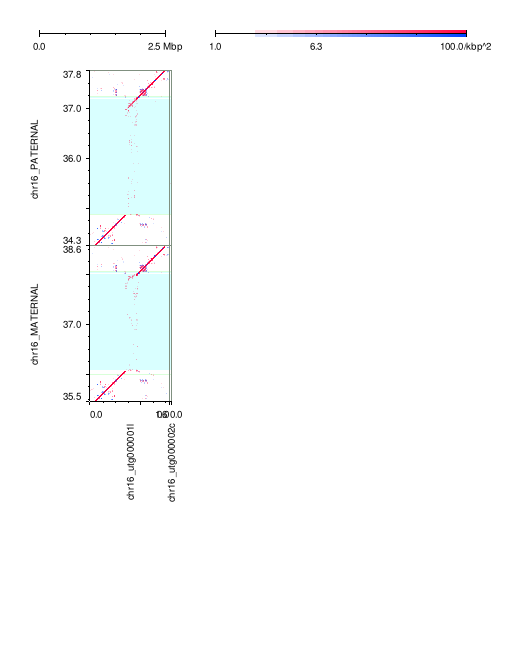

### mm2-k133-hpc.png

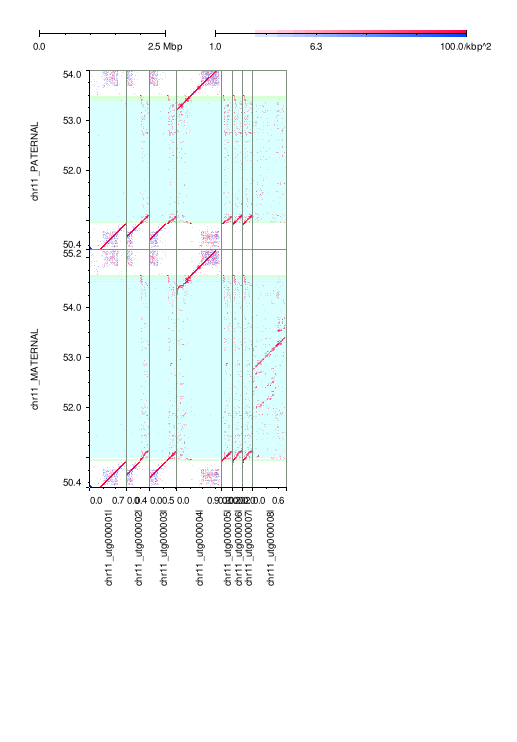

### mm2-k133-hpc.png

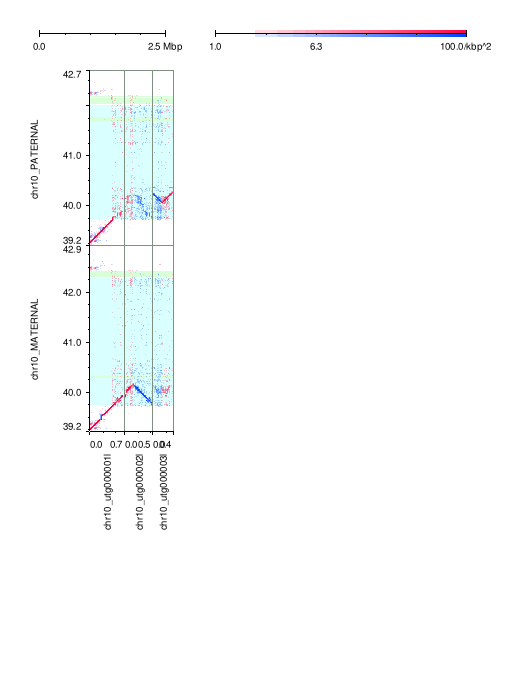

### mm2-k133-hpc.png

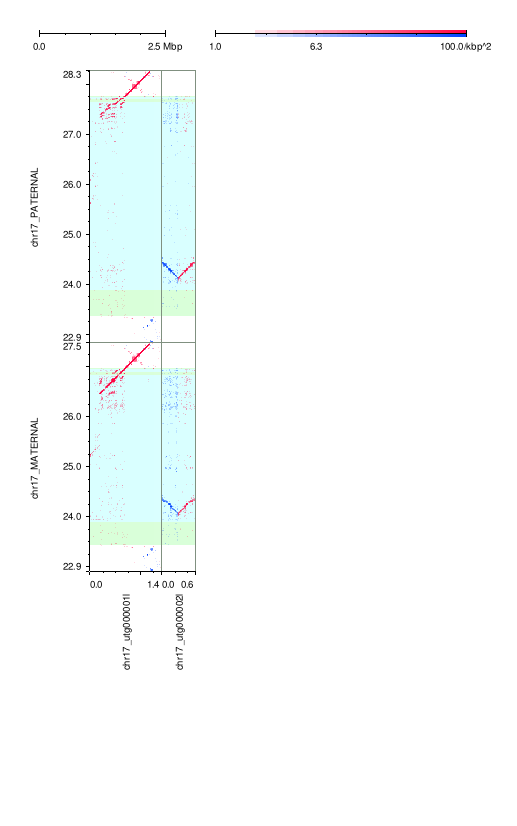

### mm2-k133-hpc.png

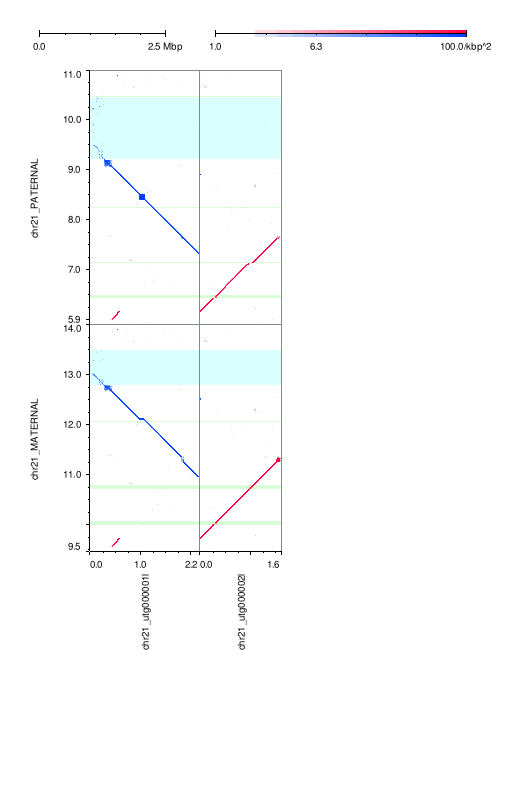

### mm2-k133-hpc.png

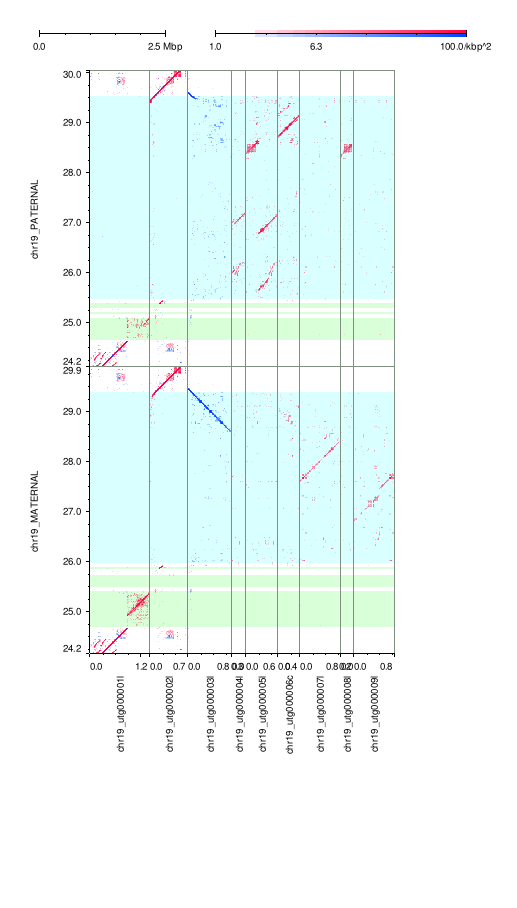

### mm2-k133-hpc.png

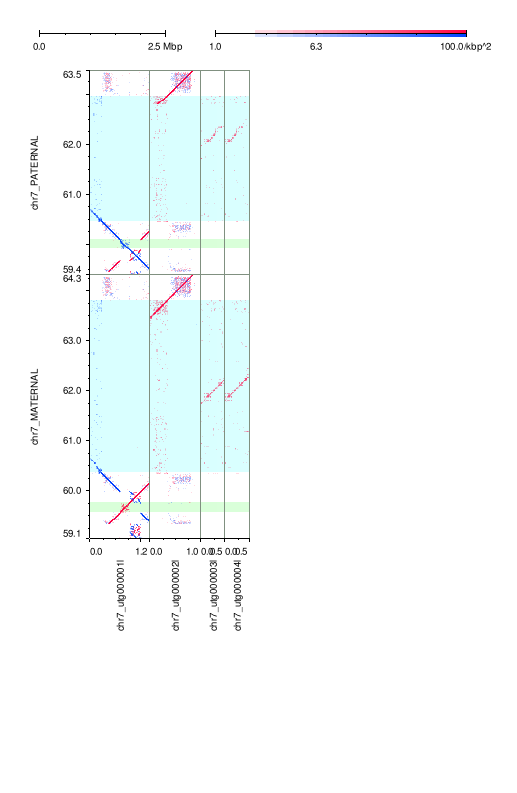

### mm2-k133-hpc.png

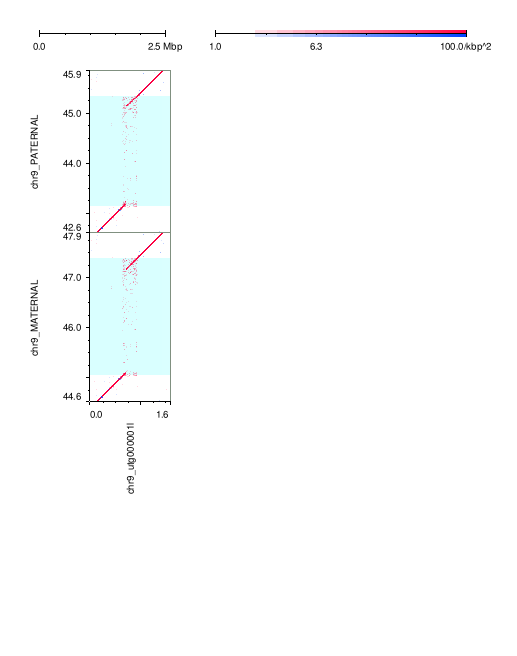

### mm2-k133-hpc.png

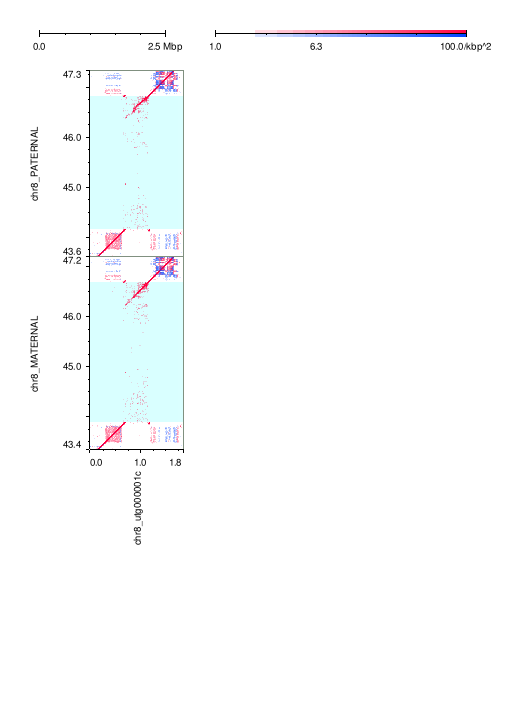

### mm2-k133-hpc.png

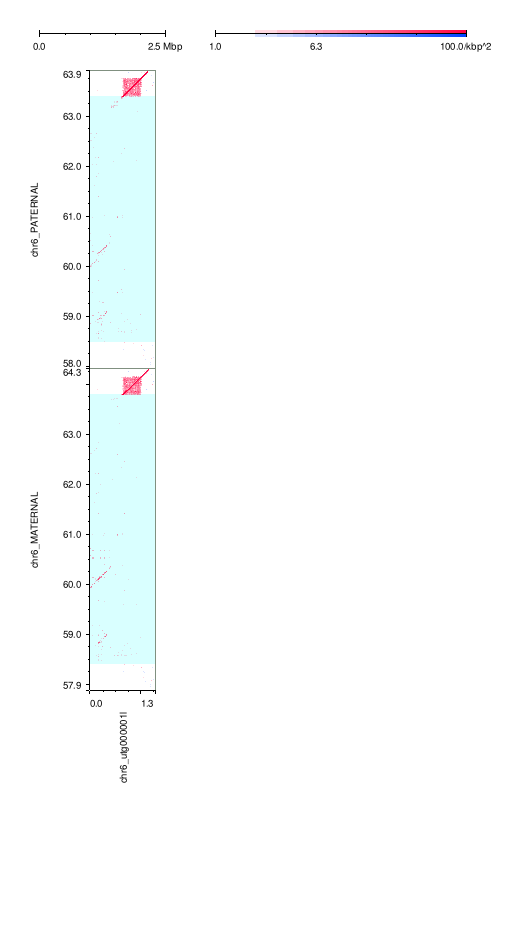

### mm2-k133-hpc.png

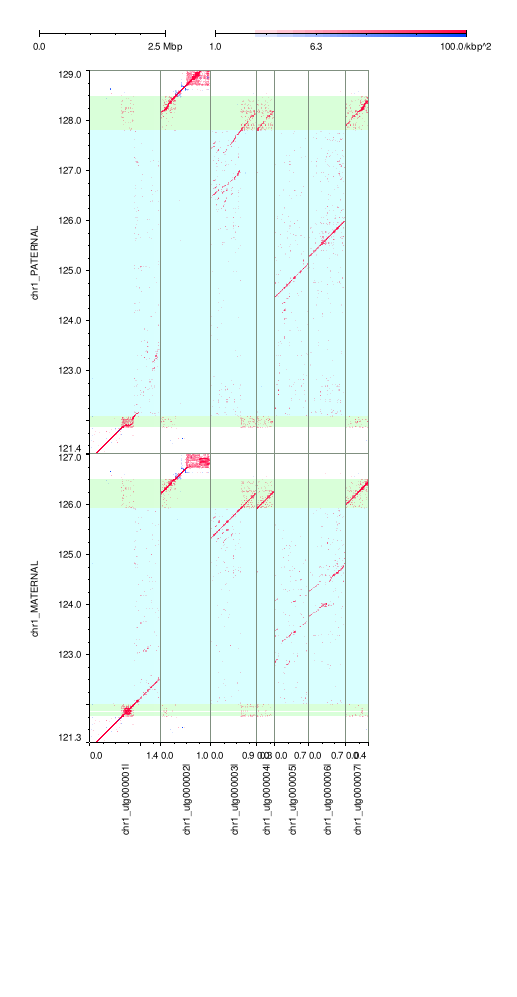
